# MOGSA: integrative single sample gene-set analysis of multiple omics data

**DOI:** 10.1101/046904

**Authors:** Chen Meng, Azfar Basunia, Bjoern Peters, Amin Moghaddas Gholami, Bernhard Kuster, Aedín C Culhane

**Author notes:** Current address: Roche Sequencing Solutions, 1301 Shoreway Road, Suite 300, Belmont, CA 94002, USA.

## Abstract

Gene set analysis (GSA) summarizes individual molecular measurements to more interpretable pathways or gene sets and has become an indispensable step in the interpretation of large scale omics data. However, GSA methods are limited to the analysis of single omics data. Here, we introduce a new computation method termed multi-omics gene set analysis (MOGSA), a multivariate single sample gene-set analysis method that integrates multiple experimental and molecular data types measured over the same set of samples. The method learns a low dimensional representation of most variant correlated features (genes, proteins, etc.) across multiple omics data sets, transforms the features onto the same scale and calculates an integrated gene set score from the most informative features in each data type. MOGSA does not require filtering data to the intersection of features (gene IDs), therefore, all molecular features, including those that lack annotation may be included in the analysis. We demonstrate that integrating multiple diverse sources of molecular data increases the power to discover subtle changes in gene-sets and may reduce the impact of unreliable information in any single data type. Using simulated data, we show that integrative analysis with MOGSA outperforms other single sample GSA methods. We applied MOGSA to three studies with experimental data. First, we used NCI60 transcriptome and proteome data to demonstrate the benefit of removing a source of noise in the omics data. Second, we discovered similarities and differences in mRNA, protein and phosphorylation profiles of induced pluripotent and embryonic stem cell lines. We demonstrate how to assess the influence of each data type or feature to a MOGSA gene set score. Finally, we report that three molecular subtypes are robustly discovered when copy number variation and mRNA profiling data of 308 bladder cancers from The Cancer Genome Atlas are integrated using MOGSA. MOGSA is available in the Bioconductor R package “mogsa”.

## Introduction

Increasing numbers of studies report comprehensive molecular profiling using multiple different experimental approaches on the same set of biological samples. These multi-omic studies can potentially yield great insights into the complex molecular machinery of biological systems. High-throughput sequencing allows quantification of global DNA variation and whole transcriptome RNA expression (1, 2). Mass spectrometry (MS)-based proteomics can identify and quantify the majority of proteins expressed in a human tissues or cell lines(3). Emerging single cell sequencing technologies enable simultaneous measurement of transcriptomes and protein markers expressed in the same cell, using CITE-seq or REAP-seq (4, 5). Integrating, interpreting and generating biological hypothesis from such complex data sets is a considerable challenge.

Gene-set analysis (GSA) is widely used in the analysis of genome scale data and is often the first step in the biological interpretation of lists of genes or proteins that are differentially expressed between phenotypically distinct groups (6). These methods use external biological information, including gene ontologies, to reduce thousands of genes or proteins into lists of gene-sets that describe cellular pathways, subcellular localization, transcription factors or miRNA targets etc., thus facilitating hypothesis generation.

Large scale omics studies or single cell studies may have limited *a priori* knowledge of phenotype groups or may aim to discover new molecular subtypes in a panel of experimental conditions or tissues with complex phenotypes, exemplified by The Cancer Genome Atlas (TCGA) (7) and the Clinical Proteomic Tumor Analysis Consortium (CPTAC) (8). Classical GSA methods which require phenotypically distinct groups (6) have limited application in such cases and several unsupervised, single sample GSA (ssGSA) methods have been developed (9–12). These methods do not require prior availability of phenotypic or clinical data. Arguably, one of the most popular approaches is single-sample GSEA that ranks genes according to the empirical cumulative distribution function and calculates a single sample-wise gene-set score by comparing the scores of genes that are inside and outside a gene-set (10). A related method, gene-set variation analysis (GSVA), also calculates sample-wise gene set enrichment as a function of the genes that are inside and outside a gene set, and also uses a similar Kolmogorov-Smirnov-like rank statistic to assess the enrichment score, but genes are ranked using a kernel estimation of a cumulative density function (9). These single-sample GSA methods are designed for the analysis of a single data set, and do not integrate or calculate a single sample GSA score on multiple data sets simultaneously.

Here, we present a novel unsupervised single-sample gene-set analysis that calculates an integrated enrichment score using all of the information in multiple omics data sets, named “multi-omics GSA” (MOGSA). The method relies on matrix factorization (MF), powerful methods that can be used to learn patterns of biological significance in high dimensional data (13) as well as identify and exclude batch effects (14). Coupled or tensor MF methods can learn latent correlated structure within and between omics data sets (15–18) and have been applied to the analysis of molecular data from different technology platforms (19) and integration of diverse multi-omics data (15, 17).

An attractive characteristic of coupled or tensor MF approaches, is that they identify global correlated patterns among samples or observations, and, therefore, do not require pre-filtering of gene identifiers in each data set to a common intersecting subset of features (genes/proteins). All features, whether they have annotation or not, can be included in the analysis, and, therefore, the method can be applied to analyzing data from experimental platforms that include unknown molecules (for example lipidomics, metabolomics), novel identifiers or molecules that are difficult to map one-to-one between data sets (e.g. transcript variants to proteins). Whilst coupled MF or latent factor methods are powerful, they identify components, the interpretation of which, can be challenging and may require domain knowledge (15, 20–22). MOGSA uses the components to calculate scores for gene sets in each biological sample, providing simple, accessible biological interpretation. We show that integrative ssGSA by MOGSA has higher sensitivity and specificity for the detection of differentially expressed gene-sets compared to popular ssGSA approaches when applied to simulated data. To demonstrate the result interpretation and application, we applied MOGSA to both small and large scale biological data from high throughput experiments.

## Experimental Procedures

### Experimental Design and Statistical Rationale

This project describes a new algorithm called MOGSA. The mathematical details of the algorithm are provided in the supplementary methods. We rigorously tested and validated the performance of the method using four distinct use-cases. In the first case, we used simulated data to benchmark the performance of MOGSA and compared it to the most widely used methods in the field. Using diverse scenarios, each with 100 simulated data sets, we demonstrated that MOGSA outperforms other methods because it can integrate multiple data sets thus increasing the sensitivity to identify gene sets with subtle perturbations. Secondly, using well-characterized multi-omics cell line data, we demonstrate the benefit of removing a source of noise (batch effect/biological bias) by excluding a component to amplify the signal in gene set analysis. We applied this to removing the effect of cell line doubling time in multiple transcriptomics data sets of 59 cell lines. The third use-case examined one of most common needs of biological laboratories; the integrative analysis of diverse molecular data obtained on a small number of biological samples. We integrated transcriptome, proteome and phosphoproteome data on four iPS ES cell lines and demonstrated how to interpret the gene set scores to reveal which data set contributed most to a specific biology process. Finally, the fourth use-case examined molecular subtype discovery using multi-omics data by the integrative analysis of copy number variation and transcriptome data of 308 bladder tumors. We demonstrate how to rigorously apply the multi-omics single sample GSA method to discover molecular subtypes and interpret the biological basis for each tumor subtype in a large scale multi-omics studies. In each case, the data are publically available, and we provide details on the methods and code such that our analysis can be reproduced.

### Data simulation

We simulated 100 multiple omics data projects. Each simulated data set was a triplet (*K*=3) containing three data matrices (Figure S1), each matrix had the dimension 1000×30, representing 30 matched observations (*n*=30) and 1,000 features (*p*_*k*_=1,000). Each data set had an annotation matrix, which assigned each feature to one of 20 non-overlapping “gene-sets”. The binary annotation matrix had dimensions of 1,000 features × 20 gene sets. Each gene set contained 50 genes. The 30 observations were defined by 6 equal sized clusters with 5 samples per cluster.

In each observation, 5 out of 20 gene-sets were simulated as differentially expressed (DE). For observations in the same cluster, the same set of DE gene-sets were randomly selected as we assumed that differentially expressed (DE) gene-sets define the difference between clusters. For a DE gene-set, a number of genes was randomly simulated as DE genes (DEG), denoted as DEG_j_. Random selection of DEGs means that the DEGs in different data sets may overlap. In separate simulations, we varied the number of DEGs per gene-set (eg 5, 10 and 25 out of 50) or mean signal to noise ratio.

We used the following linear additive model adapted from (9), the expression or abundance of gene on *i*th row and *j*th column is simulated as

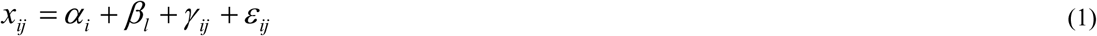

where with *i* = *1, …, p* is a gene specific effect. *β*_*l*_ ∼ N(*µ* = 0,*σ* = *s*) is the cluster effect. For observations belonging to the same cluster *l*, the same *β*_*l*_ was applied. The cluster effect factor (categorical variable) was introduced following the hypothesis that observations from the same clusters are driven by some common pathways or “gene-sets” and ensures that observations from the same cluster have a higher within than between cluster correlation. The six correlated clusters in the simulated data were captured by the first five components. The cluster effect *β*_*l*_∼ N(*µ* = 0,*σ* = *s*) was sampled from a distribution with a mean of 0 and standard deviation *s.* The standard deviation (*s*) adjusts the correlation between observations in the same cluster, and thus each cluster can have different within cluster variance and different proportions of variance would be captured by the top five components. In this study, we set s = 0.3, 0.5 and 1.0, which led to 25%, 30% and 50% of total variance captured by the top 5 components. *∈* ∼ N(*µ* = 0,*σ* = 1) is the noise factor. γ_ij_ is the differential expression factor describing if a gene is differentially expressed (DE):

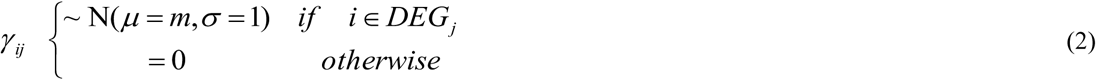

Apart from the retained variance by top five components, two other parameters were tuned in the simulation study. First, the number of DEGs in a DE gene-set (5, 10 and 25 out of 50 DEGs). The second parameter was different signal-to-noise ratios, which was tuned through modifying *m* in expression (2). The candidate values of *m* were 0.3, 0.5 and 0.8 standing for low, medium and high signal-to-noise ratio. In total, 100 projects of triplet data sets were generated. The three matrix triplets were analyzed by MOGSA. NMM, GSVA and ssGSEA, only accept one matrix as input; therefore, the three simulated matrices were concatenated into one grand matrix in these analyses. The performance was assessed by the area under the ROC curve (AUC).

### Downloading and Processing of NCI60 Cancer data

Processed mRNA expression data (normalized score averaged from 5 microarray platform) and clinical information were downloaded from CELLMINER (download date: 2017-06-01) (23). Quantitative proteome profiles were downloaded from the supplementary table of Moghaddas Gholami et al. (24). The proteome data were quantified and normalized using the iBAQ method (25) and iBAQ values were transformed by xi = log10(iBAQi + 1). The LC cell line SNB19 and melanoma cell line MDAN were missing mRNA data and were therefore excluded from the analysis.

### Determining the number of components that capture the correlated structure between NCI60 transcriptome and proteome data

A random sampling method was applied to determine the number of components that represented significant correlated structure between data sets. First, MFA was applied to NCI60 transcriptome and proteome data and the (true) variance associated with each MFA component was recorded. Next, cell line labels were randomly shuffled in both transcriptomics and proteomics data and the variance of components were calculated from the randomly labels data. We repeated this process 20 times in order to estimate the null distribution of variances associated with each component. The variance of the top three components was significantly higher than the null distribution (Figure S2).

### Downloading and processing of the iPS ES 4-plex data

The transcriptomics (RNA-sequencing), proteomics and phoshphoproteomics data were downloaded from Stem Cell-Omic Repository (Table S1, S2 and S5 from http://scor.chem.wisc.edu/data.php) (26). In this study, we used the 4-plex data, which consisted of 17,347 genes, 7,952 proteins and 10,499 sites of phosphorylation in four cell lines. For the transcriptomics data, the expression levels of genes were represented by RPKM values. Three replicates were available and we used the mean RPKM value of the three replicates. Genes with duplicated symbols and low expression values (summed RPKM < 12) were removed. The iTRAQ quantification of protein and phosphorylation sites was performed by TagQuant (27), as describe in (26). Protein and sites of phosphorylation with low intensity (summed intensity <20) were removed. In the proteomics data, proteins that were not mapped to an official gene symbol were removed. Finally, all the data were logarithm transformed (base 10). After filtering, 10,961, 5,817 and 7,912 features were retained in the transcriptomic, proteomic and phospho-proteomic data sets. A few missing values were still present and replaced with zero values. The enrichment analysis was done on the gene symbol levels, the specific phosphorylation sites were not considered.

PCA of each individual data set is shown in Figure S3. The strongest signal (first PCs) in all three data sets was the difference between NFF cells and the stem cell lines, and this difference was particularly apparent in the proteomics data sets. The second and third components represented subtle differences between iPSC and ESC lines.

### Downloading and Processing of TCGA Bladder Cancer data

Normalized mRNA gene expression, copy number variation (CNV), microRNA (miRNA) expression data and clinical information of BLCA were downloaded from TCGA (Date: 26/09/2014) using TCGA assembler (28). The processed mRNA gene expression had been obtained on the Illumina HiSeq platform and the MapSplice and RSEM algorithms had been used for the short read alignment and quantification (Referred as RNASeqV2 in TCGA) (29, 30). The gene level CNV was estimated by the mean of copy numbers of the genomic region of a gene (retrieved by TCGA assembler directly). Patients that were present in both gene expression and the CNV data were included in the analysis (n=308).

GISTIC2.0 (31) data for copy number gains/deletion in 24,776 unique genes were downloaded from TCGA firehouse (http://gdac.broadinstitute.org/; download date 2015-03-09). The GISTIC encodes homozygous deletions, heterozygous deletions, low-level gain and high-level amplifications as −2, −1, 1 and 2 respectively. The four types of events were counted for each of the patients. The total number of events was calculated by summing all four types of events

Before applying MOGSA, minimal non-specific filtering of low variance genes was performed on both data sets. RNA sequencing data (normalized count + 1) were logarithm transformed (base 10). Genes were filtered to retain those with a total row sum greater than 300 and median absolute deviation (MAD) greater than 0.1, which retained 14,692 unique genes (out of 20,531 genes). Then, RNAseq gene expression data were median centered. For the CNV data, genes with standard deviation greater than the median were retained. A PCA of each individual data set of CNV and mRNA is shown in Figure S4. From scree plots of the first 10 eigenvalues, an elbow in each plot appears between 4-6 components suggesting this number of components is needed to capture most of the variance (Figure S4), which was consistent with the reported molecular heterogeneity in these data.

### BLCA subtype clustering

To determine the optimal number of components as input to MOGSA, we performed MOGSA on the BLCA mRNA gene expression and CNV data (n=308) with a range of components from 1 to 12. For each gene-set in the GSS matrix, gene-sets were ranked by the number of tumors in which they were significantly regulated (either positive or negative GSS, p<0.05), such that gene-sets that were significant in most tumors had the highest ranks. Most gene-sets were insignificant in all tumors and no gene-set was significant in all 308 tumors. For p<0.05, we examined the 10, 20, 40, 100, 200, 500 and 1000 highest ranked gene-sets and examined the stability of gene-set ranking when additional components were included (Figure S5). Increasing the number of components (from 1 to 5) increased the stability of gene set lists, however there was little additional gain after five components in all cases (Figure S5). In addition, we confirmed that these five components were not correlated with batch effects including TCGA batch ID, plates, shipping date or tissue source sites.

Next, consensus clustering was used (32, 33) to cluster the top five latent variables with Pearson correlation distance and Ward linkage for the inner loop clustering. Eighty percent of patients were used in the re-sampling step of clustering. Stability analysis showed there was no effect when different resampling proportions (50%, 60%, 70%, 80% and 90%) were used in the inner and outer loop of consensus clustering (Figure S6). Average agglomeration clustering was used in the final linkage (linkage for consensus matrix) (32).

### Determining the number of BLCA clusters

Whilst consensus clustering analysis indicated high confidence in either two or three subtypes (Figure S7), silhouette analysis (Figure S7E) suggested three subtypes. A recent report highlighted limitations in consensus clustering (34), and therefore, in parallel, we also used the “prediction strength” algorithm, to discover the number of stable subtypes that can be predicted from the data (35). In the prediction strength method, all samples were assigned a “true” subtype label according to the clustering obtained from a given number of clusters. Then, the patients were divided into “training” and “testing” sets. The KNN classifier was used to classify the patients in the testing set. Cross-validation suggested that there is no obvious good choice of *K* (Figure S8), and, therefore, we used a wide range of odd *K* from 1-17 (Figure S9). For each test, the agreement in assignment between predicted and true labels were computed. The prediction strength was defined by the lowest proportion among all the subtypes. It indicates the similarity between the true and predicted labels and ranges from 0 to 1, where a value > 0.8 suggests a robust subtype classification (36). Therefore, the model with the greatest number of subtypes and prediction strength > 0.8 can be considered “optimal”. In this study, we performed 100 random separations of training and testing sets and the prediction strength of each randomization was calculated. The prediction strength analysis clearly supported three subtypes (Figure S9). Therefore, using two independent approaches, we determined that the data (5 components of the integrated analysis) supported three BLCA molecular subtypes.

### Sources of Gene-set annotation

Molecular Signature Database MSigDB (version 4.0) (37) gene sets included subsets C2 curated pathways, C3 motif pathways which included transcription factor target (TFT, n=617) target gene-set and C5 gene ontology (GO) biological process (BP, n=825), cellular component (CC, n=233) and molecular function (MF, n=396) terms. The pathway databases, Biocarta, KEGG and Reactome had 217, 186 and 674 gene-sets respectively. We excluded gene ontology terms that had more than 500 genes and less than 5 genes that mapped to features across all data sets. For example, in the BLCA analysis, gene-sets (1,454 in total) were filtered to exclude those with less than 5 genes in a list of the concatenated features of CNV and mRNA data resulting in 1,125 retained gene-sets.

## Results

### Outline of MOGSA algorithm

MOGSA integrates multi-omic features measured on the same set of observations (e.g. cell lines, disease tissues) and learns multi-omics patterns of gene-sets that are significantly altered in these samples (Figure 1). Omics studies can include multiple data matrices such as RNA sequencing counts of gene expression, abundance measurements of proteins, metabolites, DNA methylation, mutation or CNV data or other omics molecular data. The number of features may exceed the number of observations. We refer to genes, proteins or other biological molecules as features for simplicity below. **Figure 1** describes the three steps of the algorithm; Input omics data matrices must have matched observations but may have different and unmatched features. The number of features may exceed the number of observations. In order to map features to gene-sets, MOGSA also requires an incidence matrix of gene to gene-set membership associations for each data matrix, called “gene-set annotation matrix”, in which rows are features and columns are annotation vectors of gene-sets, a value of 1 indicates that a feature (e.g. gene) is a member of a gene-set. A feature may belong to multiple gene-sets simultaneously, that is a row sum may exceed 1.

**Figure 1:**
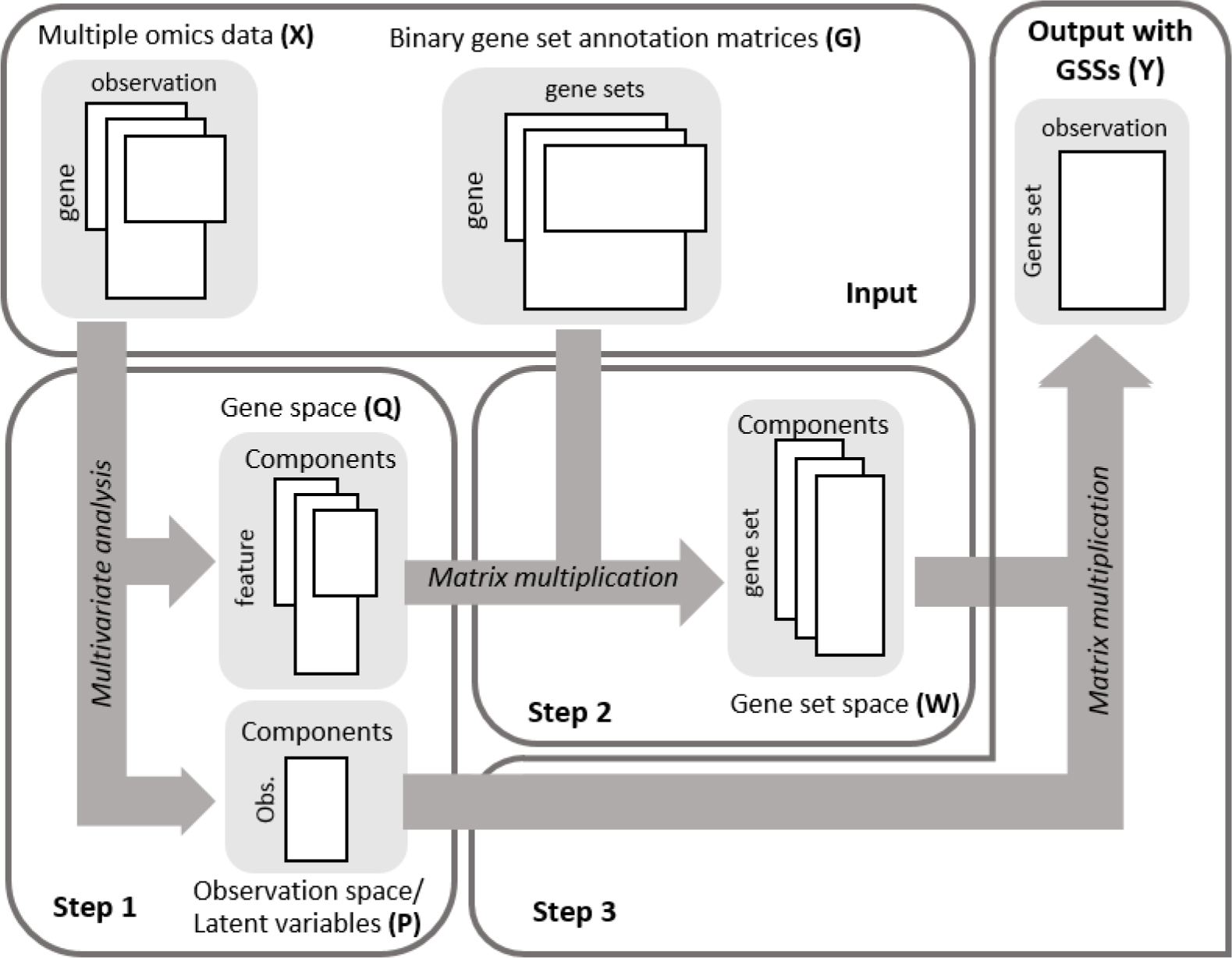
Schematic representation of the MOGSA algorithm. The algorithm requires pairs of matrices as input; multiple omics data matrices and corresponding gene-set (GS) annotation matrices. In step 1, the multiple matrices are analyzed using a multivariate analysis (MVA) method resulting in an observation space and gene space. Next, the gene-set annotation matrices are projected on the same space, and the resulting matrix contains the gene-set space. The last step is to reconstruct gene-set-observation by multiplying the observation and gene-set spaces.

In the first step, several (*k*) input data matrices are integrated using multiple factor analysis (MFA) (20). MFA is a multiple table extension of principal component analysis (PCA) that is well suited to integrating multiple omics data since it reduces high dimensional omics data to a relative small number of components that capture the most prominent correlated structure among different data sets (20, 38). To prevent data sets with more features or different scales dominating a MFA result, each data set is divided by the first eigenvalue of a decomposition of each individual data set. MFA generates matrices of latent variables (components) in observation (**P**) and feature (**Q**) space (Figure 1). The number of components is typically less than the number of observations. We retain and examine the first few components as these represent most of the variance in the data. One can select non-contiguous components (to filter or omit one or more component). Approaches for choosing the number of components are discussed later. MOGSA makes use of an attractive property of MF approaches in that supplementary data such as gene-set information (e.g. Gene Ontology annotations) can be projected onto the observations space to aid interpretation (16, 17, 39). In the next step (step 2, Figure 1) each gene-set annotation matrix (**G_1..k_**) is projected as additional information onto the gene-set space (**Q_1..k_**) generating a score for each gene-set in the same projected space (**W_1..k_**), where *k* is the number of data sets. In the final step (step 3, Figure 1), MOGSA multiplies the latent variables of the observations (**P**) and latent variables of gene-sets (**W_1..k_**) to generate a matrix (**Y**) with a gene-set score (GSS) for each gene-set in each observation (**Y**).

A gene-set with a high GSS are driven by features that explain a large proportion of the global correlated information among data matrices. These features could be from one, many or all data matrices, and may be non-overlapping. For example a GSS of a gene set with features A-H, could be driven by high levels of gene expression in genes A,B,C, and increased protein levels in proteins C,D,E and amplifications in copy number in gene H. The GSS matrix (**Y**) may be decomposed with respect to each data set (**X**) or latent variable space (**P**,**Q**) so that the contribution of each individual data set or component to the overall score can be evaluated (see Experimental Procedures for more details).

### Multiple omics GSA has greater sensitivity to identify regulated gene-sets

Biological pathways are regulated by a complex network of cellular molecules that can be measured using transcriptomics, phosphoproteomic, metabolomics, lipidomics or other molecular profiling methods. Popular ssGSA approaches, such as ssGSEA and ssGSVA are designed for analysis of a single data set. We postulated that there would be a greater power to detect gene sets or cellular pathways if methods integrated more diverse data types. To this end, we compared the performance of MOGSA to ssGSA methods that were developed for the analysis of one data set including the widely used GSVA and ssGSEA and naïve matrix multiplication (NMM) methods (9, 10).

Figure 2 shows the performance of each method applied to 100 simulated data sets. In each data set, we simulated a study of 30 observations with three omics data types that measured 1,000 features each (Figure S1; see Experimental Procedures section). Each features was a member of one of the 20 gene-sets. Each gene-set had 50 genes. The observations were grouped into 6 clusters and each clusters had 5 differentially expressed (DE) gene-sets compared to the other observations. The triplets were analyzed by MOGSA directly, however matrices were concatenated for NMM, GSVA and ssGSEA as these methods can only accept one matrix as input.

**Figure 2:**
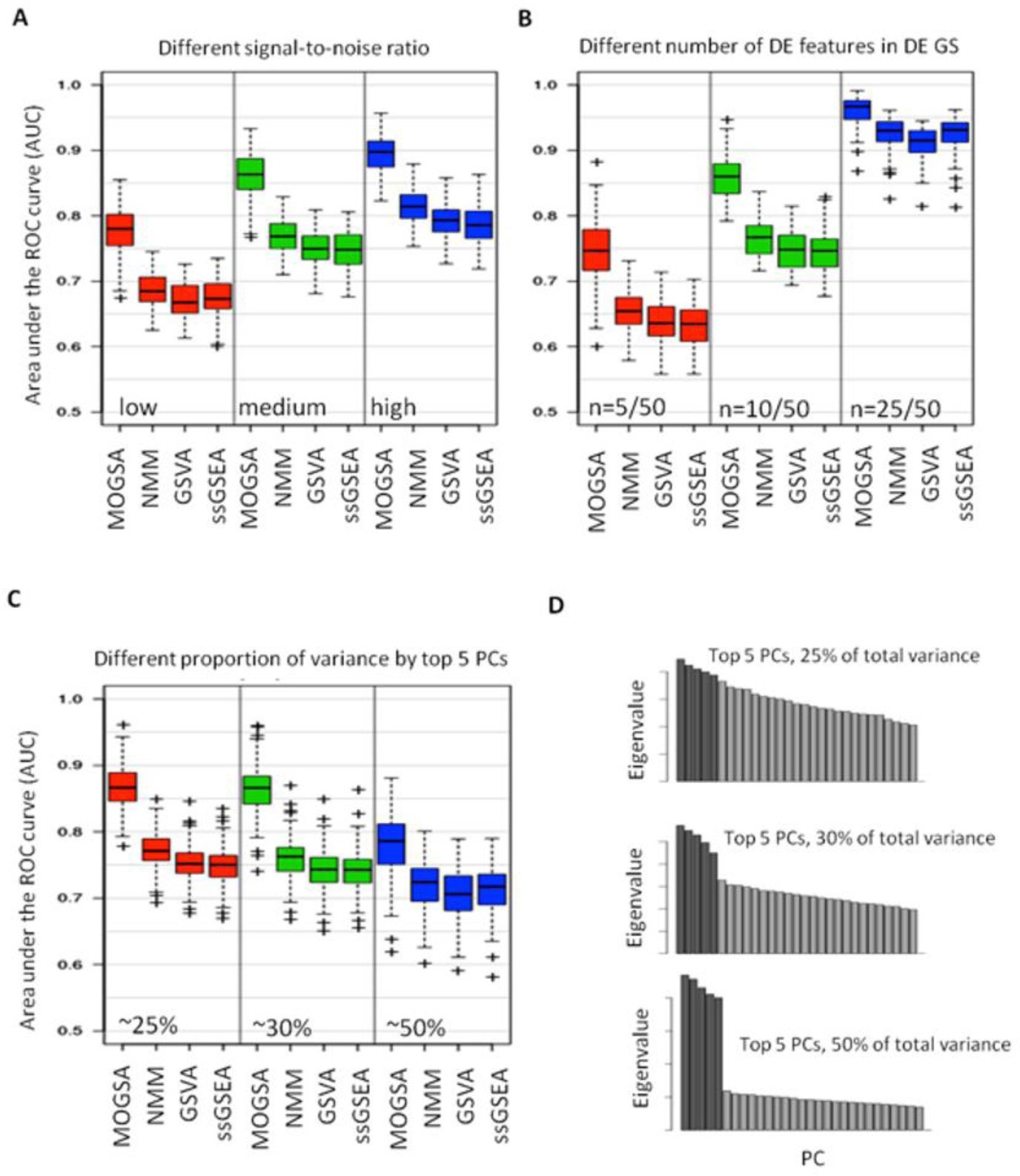
Comparison of MOGSA with NMM, GSVA and ssGSEA. The performance of each method was accessed by their ability to identify differentially expressed gene-sets over 100 simulations in every condition (as indicated by the area under the ROC curve; AUC). (A) Comparison of GSA methods using data with different signal-to-noise ratios. (B) Comparison of data with different number of differentially expressed (DE) genes in each of the DE gene-set. From left to right, 5, 10 and 25 of total 50 genes are differentially expressed in each of the three simulated data matrices if a gene-set is defined as DE gene-sets. (C) Scree plots show representative eigenvalues in each of the conditions in (D). (D) AUCs with different proportion of variance are capture by the top 5 components. From left to right, 25%, 30% and 50% of total variance are captured. The darker bars represent the top 5 components.

We expected that MOGSA would be especially powerful at identifying altered gene-sets in heterogeneous noisy data. That is because MOGSA, uses only the top few most informative latent variables, thus omitting the signal of many features with little variance, which are potentially noise. Therefore we explored the power of the methods to detect DE gene-sets when there was a strong or weak gene expression signal. First we simulated increasing DEG signal-to-noise by changing the mean gene expression of DEGs in the cluster and secondly we altered the number of DE genes in a DE gene-set (5, 10 and 25 genes). Specificity and sensitivity of the methods detecting the DE gene-sets were evaluated and the performances were summarized as the area under the receiver operating characteristic curve (AUC). As expected, the performance of all methods was better when the signal-to-noise ratio or the number of DE genes in DE gene-sets increased (Figure 2A and 2B). MOGSA consistently outperformed the other methods and the difference was even more apparent when the signal-to-noise ratio was low or when there were few DE genes (5 or 10 of 50 genes) (Figure 2B).

Next we compared the performance of each method using data with a simple or complex structure which reflect the complexity of phenotypes in a study. In data with a simple phenotype, a few components should easily capture most of the variances. However in data with a complex phenotype, for example a heterogeneous tumor data set, with mixed histology, grade and response to treatment, there are many signals so that many latent variables may be required to capture even half of the variance. In the simulated data, observations grouped into six clusters, which could be captured by the first five components. Therefore, we simulated data such that the first 5 components captured 50%, 30% or only 25% of the total variance (Figure 2C). Again, MOGSA outperformed the other methods and was relatively robust to changes in the variance retained (Figure 2D). The performance (AUC) of all methods decreased when greater variance was retained, which can be explained by higher intra-cluster correlation that leads to a lower signal-to-noise ratio in data sets (see Experimental Procedures).

Given the many fundamental differences between MOGSA and the other ssGSA methods, we repeated the simulations adjusting for technical aspects of the MOGSA approach that might give it an “unfair edge”, but these did little to improve the performance of the other methods. Because GSVA and ssGSEA were designed for analysis of single data sets, we compared the performance of GSVA and ssGSEA on a single data sets of the triplet compared to the concatenated triplet. Concatenating multiple data matrices neither improved nor decreased the performance compared to analysis of single data sets, which is most likely caused by the signal-to-noise ratio increased accordingly with concatenation (Figure S10). In addition, since MFA weighs input matrices by their first singular value before MOGSA, we examined the effect of data set weighting on the other methods and found MOGSA still outperformed ssGSEA and GSVA when data matrices of the triplet were weighted before concatenation (Figure S11).

### MOGSA provides an easy-approach to selectively filter or enrich for components

Large scale omics studies often have several sources of unwanted variance, due to technical batch effects (40) or unwanted biological sources, for example, tumor purity in bulk tumor studies (41) or cell cycle genes in single cell molecular studies (42). In our previous study, that integrated the proteome and transcriptome of 60 National Cancer Institute (NCI) cell lines, we observed that one of the top components in both data sets was correlated with cell doubling time (24). Cell doubling time in cell culture is a characteristic of the cell culture conditions and may not reflect the cell doubling time when cells are grown in more physiologically relevant three dimensional culture models (43) or indeed those of tumors *in vivo*. We hypothesized that MOGSA would capture the doubling time effect as a component. A benefit of MF methods is that we can specifically exclude a component in order to remove the variance associated with it. In this case, by removing the component associated with doubling time (a feature of *in vitro* cell culture) we can potentially observe different gene set scores which might be closer to *in vivo* biology. In our analysis, the significantly correlated structure between transcriptomic and proteomic data was captured in three components (see Figure S2 and Experimental Procedure) and the first component which represented 6% variance was significantly correlated with cell doubling time (Figure S12, Pearson Correlation R = 0.61, p=3×10^−7^). Excluding the first component from score computations, removed the effect of cell doubling time (Figure S13), generating alternative gene set scores. For example, the gene set score (GSS) for cell cycle checkpoint was significant (p<0.001) in 31 cell lines, and after excluding the first component, was no significant in any of the cell lines (Figure 3A).

**Figure 3:**
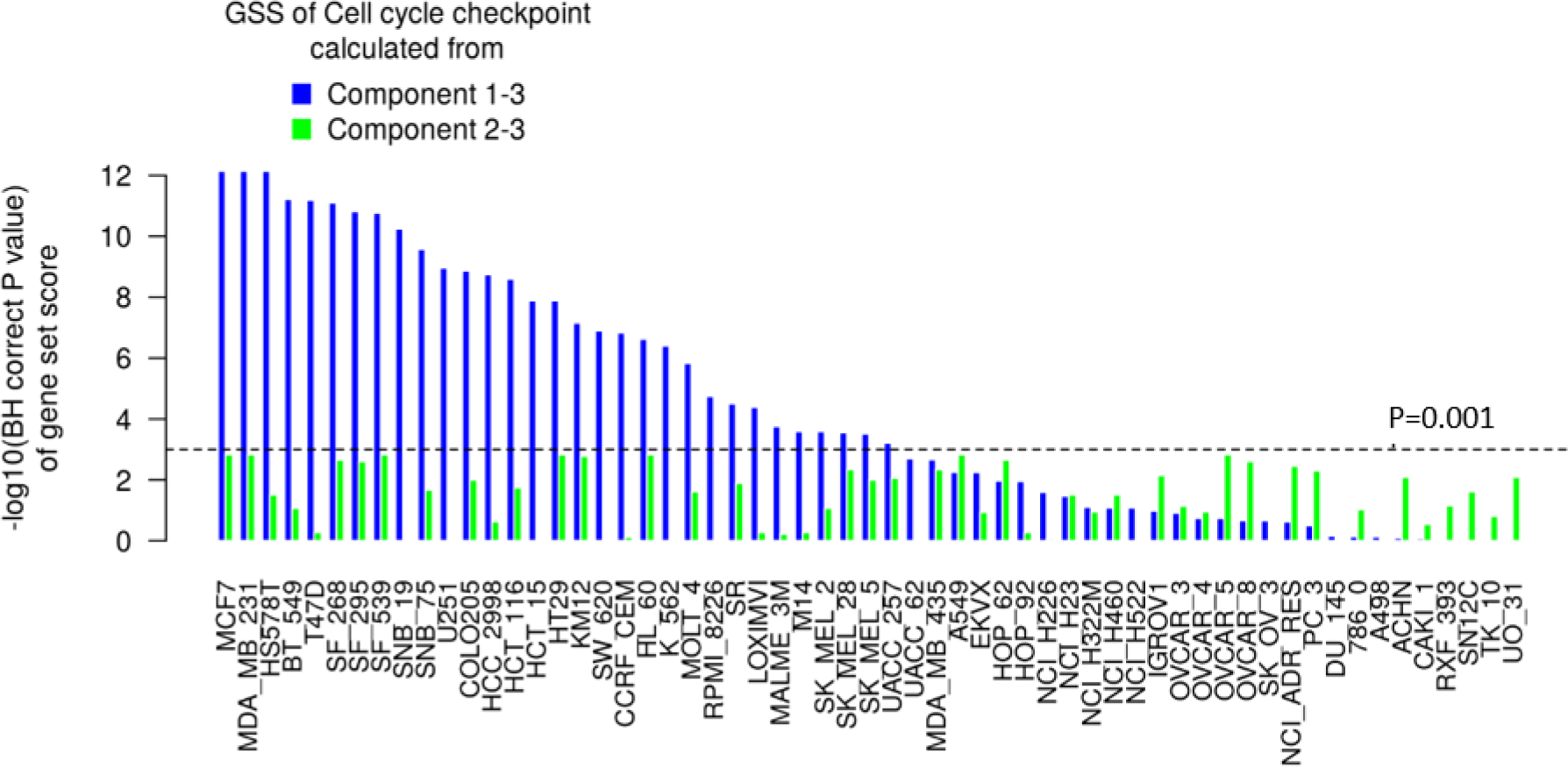
Different gene set scores by excluding components corresponding to unwanted variance. (A) The BH corrected P value of cell cycle checkpoint pathway (REACTOME) is increased by excluding components well correlated with cell doubling time (component 1). The corrected p value increased from <10^−12^ to >10^−3^ in MCF7 and MDA_MB_231 cells because component 1 is the driving force of statistical significance in these two cell lines (see Figure S13).

### Application of MOGSA to the interpretation of stem cell mRNA and proteomics data

We applied MOGSA to study profiles of mRNA (n=10,961), protein (n=5,817) and phosphorylation sites of proteins (n=7,912) of four cell lines - two embryonic stem cell lines (ESC; H1 and H9), one induced pluripotent cell line (iPSC; DF19.7) and a fibroblast cell line (newborn foreskin fibroblast; NFF). Induced pluripotent stem cells (iPSC) are adult cells that have been reprogrammed to be more like embryonic stem cells (ESC) and have great potential in the field of regenerative medicine. These cells express ESC markers and can differentiate into different cell types (26). Induced pluripotent cells are often derived from NFF cells.

MFA recapitulated the PCA of the individual data sets (Figure S14, S3). The three data sets contributed similar variance in the integrated analysis, as indicated by the weights of each data set in MFA. The variance of the first PCs were 0.24, 0.26 and 0.26 for the transcriptome, proteome and phospho-proteome data set respectively. Most of the variance was captured in the first component and it discriminated between NFF and other cell lines. The variance of the molecular differences between the ESC cells (captured by the second component) was greater than the difference between ESC and iPSC cell lines (component 3) (Figure S14).

MOGSA of gene ontology (GO) biological processes (BP) discovered 228 GO terms (out of 825) that had significant up or down-regulated gene-set scores (GSSs) in at least one cell line (BH corrected p value < 0.01). These 288 GO terms grouped into 21 broad categories, when overlaps of gene membership to GO terms was reduced using hierarchical clustering analysis (Hamming distance and complete linkage) (Table S1). GSS of representative GO terms from each category are shown in Figure 4A. Biological processes associated with more differentiated cell types were associated with the NFF cells and included up-regulation of vesicle-mediated transport, immune related responses and cell adhesion. In contrast, cell proliferation GO terms such DNA replication, and cell cycle processes had significantly higher GGS in the highly proliferative stem cell lines. These results are consistent with previous findings (26).

**Figure 4:**
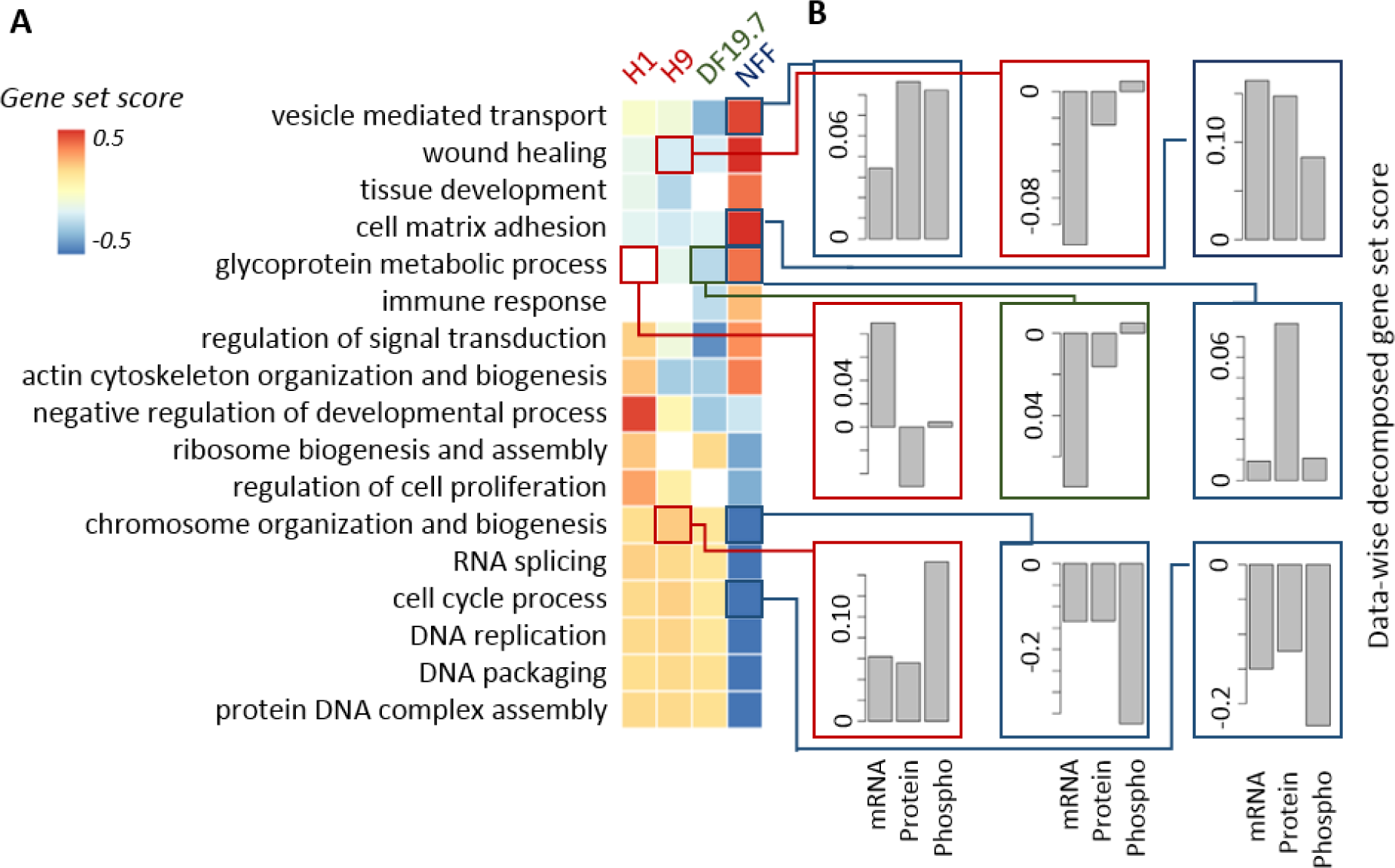
integrative gene-set analysis of iPS ES 4-plex data. (A) Heatmap showing the gene-set score (GSS) for significantly regulated gene-sets in the cell lines. The white colored blocks/cells indicate that the change of gene-sets are non-significant. (B) Data-wise decomposition of the GSS for some of the gene-sets. The contribution of each of the data is represent by a bar. The Y-axis is the data-wise decomposed gene-set score.

In the integrative analysis of multiple omics data, it is important to evaluate the relative contribution (either concordant or discrepant) of each data set to the overall GSS. Data-wise decomposition of the GSSs (see supplementary methods) are shown in Figure 4B. The three data sets had concordant contributions to most of the GO terms, including vesicle mediate transport, cell matrix adhesion, cell cycle processes in NFF line; chromosome organization and biogenesis in H9 and NFF cell lines. However, we also observed differences in the contribution of mRNA, proteins and phosphoprotein data to the GSS, for example chromosome organization and biogenesis had significant positive GSS in the stem cells and significant negative GSS in the NFF cells, and were driven by differences in the phosphorylation data. Another case where the mRNA and protein were incongruent is the GO class “glycoprotein metabolic process”. The GSSs of this term are 9.7 (p<0.001), −8.6 (p<0.01), −5.3 (p<0.01) and 0 (p>0.05) in NFF, iPSC, H9 and H1 cells respectively. Up-regulation in NFF mainly reflected up-regulation on the protein level. However, down-regulation in iPSC DF19.7 cells was due to low expression of related mRNAs. The GO term “wound healing” has previously been shown to be differentially upregulated in fibroblast NFF cells compared to ESC (26). Consistently, we also found wound healing to be upregulated in NFF compared to ESC; the GSS for wound healing were 14.2 (p<0.01), −5.4 (p<0.01), −5.2 (p<0.01) and −3.6 (p<0.001) for NFF, iPSC, H9 and H1 cells respectively (Table S1). Down-regulation of wound healing in H9 cell line was dominated by mRNA data, and the two proteomics data sets contributed little to the negative GSS. In contrast to previous studies (26), we did not observe significant differences in wound healing between iPSC and ESC. This difference could be because MOGSA is more sensitive (than single data GSA) in detecting gene-sets that have subtle but consistent changes in multiple data sets. More importantly, the contribution of individual gene-sets could be evaluated by the decomposition of GSS with respect to data sets

### Application of MOGSA to molecular subtype discovery in bladder cancer

BLCA is a molecularly heterogeneous cancer with between 2 and 5 molecular subtypes (reviewed by (44, 45)). Briefly, Sjödahl *et al.* first defined five major subtypes termed urobasal A (UroA), UroB, genomically unstable (GU), squamous cell carcinoma-like (SCCL) and ‘infiltrated’ (36). The TCGA study defined four expression clusters (I–IV) (44). The two subtype model consists of basal-like and luminal subtypes (46) which was extended by Choi et al. who defined a ‘p53-like’ luminal subtype apart from basal-like and luminal subtypes (47). Since MOGSA performs unsupervised integrative ssGSA, the integrative components can be applied to cluster discovery in multi-omics data (48) and the incorporation of gene set annotation will greatly facilitate the interpretation of the defined subtypes. We learned an integrative subtype model of BLCA in an MOGSA of CNV (n=12,447) and mRNA (n=14,719) data of 308 muscle invasive urothelial bladder cancer (BLCA) patients (obtained as part of the TCGA project). In our analysis, the top five MFA components captured a quarter of the total variance and were not dominated by either CNV or mRNA (CNV 50.6%, mRNA 49.4%; Figure 5A). We performed extensive analyses to confirm that 5 components captured sufficient variance and was the optimal number of components as input to MOGSA (see Experimental procedures). We examined 1,125 gene-sets using MOGSA in every individual patient. The number of significant MOGSA gene-sets per patient (p<0.05) ranged from 183 to 595 (Figure S15). Each patient had both positive (up) and negative gene set scores (GSSs). We focused on the 73 gene-sets that were significantly regulated (positive or negative GSS scores, p value<0.05) in most patients (≥ 200 of the 308 patients) (Table S3 and Figure S16). Cluster analysis of the GSS matrix (73 selected gene-sets x 308 tumors) revealed three clusters of gene-sets.

**Figure 5:**
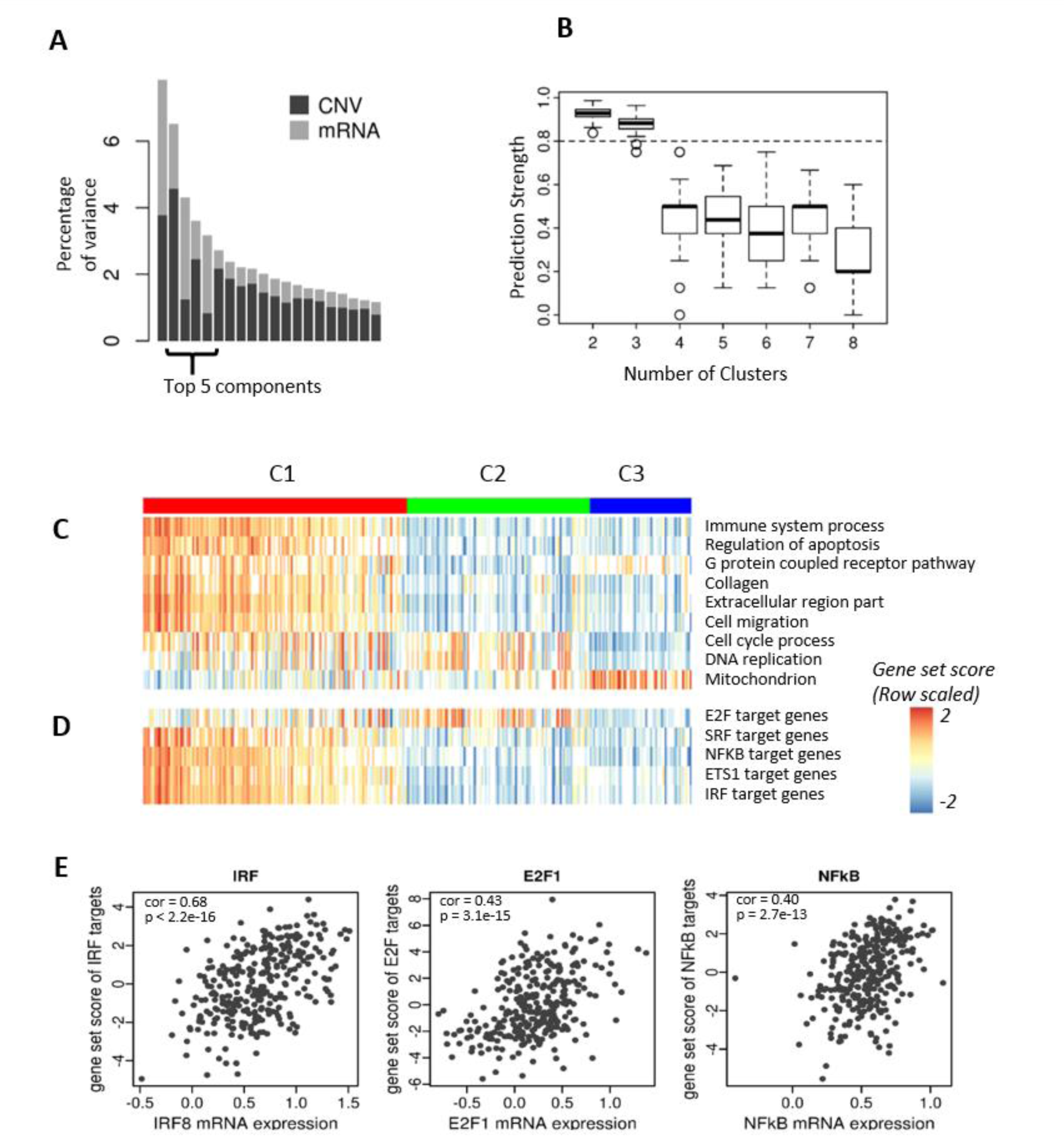
Data integration using MOGSA and integrative subtype defined by latent variables. (A) Bar plot showing the eigenvalues of components defined by MFA. The top 5 components were selected in the analysis. (B) Prediction strength was used to evaluate the robustness of classification into two to eight subtypes. The boxplot shows the prediction strength of 100 randomizations. Two and three are relative robust subtype models (prediction strength > 0.8). (C) Gene ontology (GO) and transcriptional target (TFT) gene-sets annotation of tumors. Heatmap showing the GSSs for selected gene-sets. The gene-sets “immune-related, apoptosis, G protein receptor, collagen, extracellular region and cell migration” are strong in the C1 (basal-like) subtype, whereas the mitochondrial related gene-sets are over represented in the C3 (luminal A-like) subtype of tumors. (D) The most significant transcriptional factor (TF) target gene-sets. The gene-set scores suggest that 4 out of the 5 TFs are hyperactive in the C1 subtype, except E2F family is active in the C2 subtype of cancer. The white spaces in (A) and (B) denote non-significant GSSs. (E) The scatter plots display the correlation between gene-set scores and the mRNA level of selected TFs. The expression of selected TFs is significantly correlated with their gene-set scores (also see Figure S21).

### Validation of the number of BLCA clusters

In order to identify the number of robust BLCA clusters, we applied several cluster analyses to the five components (which were input to MOGSA) and tested how many clusters were supported by the data. We applied rigorous cluster robustness analysis using methods including prediction strength and confirmed that the data supported three subtypes (Figure 5B), the final three subtype model were learned from consensus clustering (32) of the five components and the robustness of the three BLCA subtype model was extensively validated (see Experimental procedures). The three integrative BLCA subtypes consisted of two larger subtypes C1, C2 containing 148 and 103 patients respectively, and a robust smaller group C3 with 57 patients (Figure 5C).

The clusters obtained by hierarchical clustering analysis of the 73 gene-sets scores from MOGSA were consistent with the three clusters obtained by consensus clustering of the five components. A large number of GSS were associated with the C1 patients (p<0.05), more than C2 or C3 (Figure S16). A large cluster of 51 gene-sets had positive GSS scores in C1 but negative scores in C2 or C3. Two smaller clusters of gene-sets of 16 and 6 gene-sets had positive GSS scores in C2 and C3 respectively (Figure S16).

We compared the overlap between the integrative BLCA subtypes and with BLCA subtypes reported in previous studies (44) (Table S2, Figure S17). Subtypes C2 and C3 were similar to the Damrauer luminal subtype (46). But, the C3 subtype contained more low-grade tumors and showed a strong overlap with the UroA subtype of the Sjödahl study and type I of the TCGA subtype model. Subtype C2 tumors overlapped with the genomically unstable subtype defined by Sjödahl (Table S2). Accordingly, we observed higher mutation rates in the C2 patients (Figure S18). The integrative subtype C1 included a high number of patients in the type III and IV of the TCGA subtypes, the infiltrated and SCCL subtypes of the Sjödahl study (36) and the basal-like subtype identified by Damrauer (BH corrected p-value < 0.05, Table S2) (46).

### Gene sets enriched in the 3 BLCA molecular subtypes

The large number of the C1 basal-like/SCC-like BLCA gene-sets (31/51) were associated with “immune response”, which supports associations between immune regulation and the basal-like cluster that have been previously reported (36) The most significant gene-sets, “immune response” and “immune system process” had significant positive or negative GSS in 270 and 265 of 308 patients respectively (Table S3). The median GSS for the gene-set “immune system process” was 0.82, −0.75, −0.61 in C1, C2 and C3 respectively (Figure 5C;S19) indicating that immune related processes have high gene expression or copy number amplification in the C1 subtype and much lower in C2 and C3.

The remaining 20/51 gene-sets in the C1 cluster of gene-sets reflected “extracellular”, function, cell morphogenesis, migration and muscle cell development, “apoptosis” (2 gene-sets), and “G protein coupled receptor” (6 gene-sets) (Figure 5C;S19) and EMT related gene sets (Figure S20), which is consistent with reports that the Basal-like subtype has more muscle-invasive and metastatic disease at presentation (36). Gene-sets related to “cell cycle” (9 gene-sets) and “DNA repair and chromosome related” (7 gene-sets) had high GSS in luminal gnomically unstable C2 (49) (and some C1) and “mitochondrion” (4 gene-sets) in C3 (Figure 5C;S19). The mitochondrial component has been described in bladder cancer and other cancers previously (49, 50) and our study particularly associated this function with C3 low-grade papillary-like subtype in BLCA.

### Discovery of transcription factors that might regulate gene expression in BLCA subtypes

To identify transcription factors (TF) that may regulate gene expression in the three tumor subtypes, we used transcriptional factor target (TFT) gene-sets to annotate the tumors. Similar to the selection of GO terms, we focused on TFT gene-sets with more than 200 significant GSSs across 308 patients (Table S2). The GSSs of the E2F family target gene-set were significantly different in most of the tumors and are particularly low for the C3 tumors. Four identified TFs that were highly elevated in the C1 subtype; *MADS* (*MCM1*, Agamous, Deficiens, and *SRF*) box superfamily member, *SRF* and several TFs associated with transactivation of cytokine and chemokine genes, including *NFkB1, ETS1* and *IRF1* (Figure 5D). The genes exhibiting the largest GIS in the *IRF1* and *NFkB1* target gene-sets included *ACTN1, CXorf21, ICAM1, MSN, TNFSF13B, IL12RB1* and *CDK6* (Table S5). Further, we examined the correlations between GSSs and the mRNA expression. All five TFs showed that the TF mRNA and GSSs are significantly correlated (Figure 5E, Figure S21). The boxplot of GSS distribution in different are shown in Figure S19.

### The gene influence score (GIS scores)-evaluating influence of individual genes in gene set scores

In studies trying to define molecular biomarkers, a single or small number of genes or proteins are often preferred as opposed to a gene set, which may contain hundreds of genes. To evaluate the importance of individual features in a gene-set, we calculated a gene influential score (GIS) using a leave-one-out procedure (see supplementary methods). The maximum GIS value for a gene in a gene-set is 1, which indicates that this gene contributes a high proportion of variance to the overall variance of the GSSs. A GIS close to 1 often suggests a high correlation between the gene expression value and GSS. Gene influential scores of the gene-set immune system process in BLCA suggested that the top ranked genes included *ITGB2, SPI1, DOCK2, LILRB2* and *LAT2*. Other highly ranked genes included drug target genes such as *CD4, IL6*, the interferon induced proteins *IFITM2* and *IFITM3* and the G protein coupled receptors *GPR183* and *CMKLR1* (Table S4). Moreover, the C3 subtype tumors had higher GSSs in mitochondrial related gene-sets and lower expression of genes related to cell cycle process and DNA replication. GIS analysis suggested that two families of genes, NADH dehydrogenases (NDUFs) and mitochondrial ribosomal proteins (*ABCC1*/*MRP*) influenced the mitochondrial proteins (Table S4).

### Decomposing GSS scores by data-type

We decomposed the GSSs with respect to the data sets and performed a gene influential score (GIS) analysis to learn the mRNA and CNV features that influenced each GSS. For example, Figure 6A shows a GIS analysis of the GSS “cell cycle process”, where we found that mRNA expression strongly influenced the GSS, particularly the low GSS of the C3 subtype patients. The top 30 most influential genes were all observed in mRNA expression data (Figure 6B), including *RACGAP1, DLGAP5, FBXO5, AURKA, KERA* (*CNA2*) and *CDKN3* (Figure 6C). Indeed, data-wise decomposition of GSS identified several GSSs that were driven by the mRNA data, including the immune system process, DNA replication and mitochondrion gene-set (Figure S22). By contrast, both CNV and mRNA data influenced the gene-set “G protein coupled receptor activity” (Figure 6D) and the most influential genes (the GIS analysis) included both mRNA and CNV data (Figure 6E), and these differentiated the C3 subtype (Figure 6F). The features that scored highly in the GSS “G protein couple receptor activity” included CNV of *GRM6, NMUR2* and adrenergic receptors, and the gene expression of *ADGRL4* (*ELTD1*), *CMKLR1* (Figure 6F). In this gene set, the cancer driver gene and drug target *PDGFRB* were identified on both CNV and mRNA levels.

**Figure 6:**
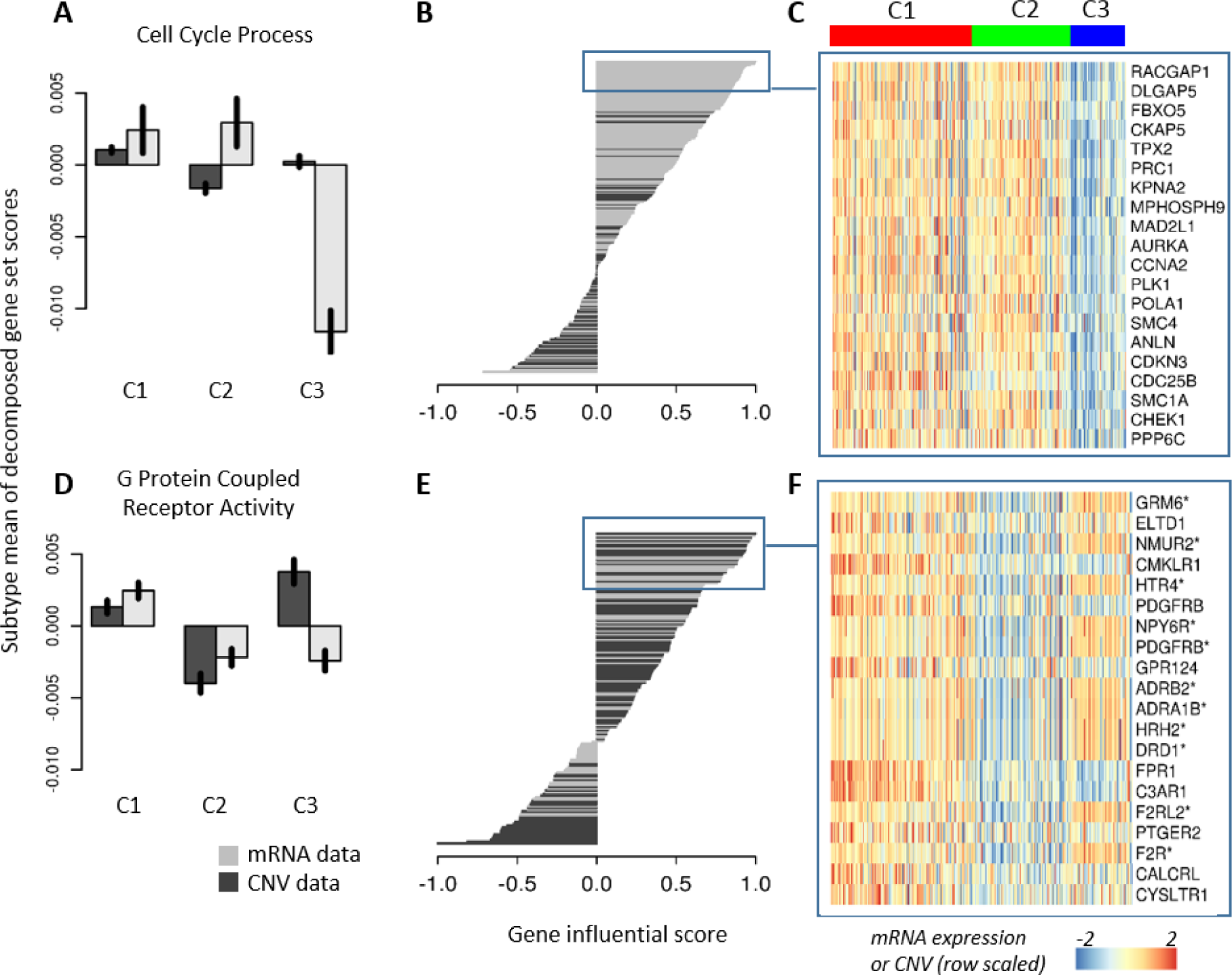
CNV and mRNA data contribute unequally to defining subtype and gene-set scores. (A) Data-wise decomposition of gene-set scores for “cell cycle process”. The bar plot shows the normalized mean of data-wise decomposed GSSs in each subtype (the black vertical line on the bars show the 95% confidence interval of the mean). (B) The bar plot shows the gene influential scores (GISs) of genes in the “cell cycle process” gene-sets. The expression of the top 30 most influential genes in the gene-set are shown in (C). (D-F) Same as (A-C) for “G protein couple receptor activity”. Gene names in (F) with asterisks indicate genes from CNV data.

### Computational performance

We applied MOGSA to various data set sizes and compared its computational efficiency to other methods. It performs comparably or marginally slower than ssGSEA, but it is faster than GSVA (Figure S23) and much faster than coGAPS (data not shown as it failed to compute on larger breast cancer and panCancer TCGA data sets in our studies).

## Discussion

In this manuscript, we introduced a new approach for multi-omics ssGSA, termed MOGSA that enables the discovery of e. g. biological pathways with correlated profiles across multiple complex data sets. MOGSA uses tensor MF or multivariate latent variable analysis to explore correlated global variance structure across data sets and then extracts the gene-sets or pathways with the highest variance that are most strongly associated with this correlated structure across observations. By combining multiple data types, we can compensate for missing or unreliable information in any single data type so we may find gene-sets that cannot be detected by single omics data analysis alone (15).

MOGSA is fundamentally different from other gene-set enrichment analysis methods which use a ‘within observation summarization’ such as the mean or median of gene expression of genes in a gene-set (9, 10, 38). MOGSA has several unique characteristics that make it well suited for generating integrated multi-omics gene set scores. First, MOGSA uses MFA, a multi-table extension of PCA to reduce the complexity of the original data by transforming high dimensional data to a low dimensional representation of the data on a few orthogonal components (latent variables). The components with the highest eigenvalues capture the most prominent or variant structure that is shared among the different data sets. Keeping the first few components (with high eigenvalues) and excluding lower ranked components with low variance may increase the signal-to-noise ratio of data, as they may be associated with low variant, noise or artifact variance (51, 52). In MOGSA, the entire set of features from each platform is decomposed onto a lower dimension space. The linear combination of feature loadings is used in the calculation of the gene-set scores. Features that contribute low variance contribute little to the score and thus the dimension reduction within MOGSA provides an intrinsic filtering of noise. The advantages of intrinsic variance filtering of features can be clearly seen when we applied MOGSA to simulated data. MOGSA outperformed ssGSA approaches including ssGSEA and GSVA which do not include a noise-filtering component. Second, data integration of features is achieved at the gene-sets level rather than scoring individual features. This greatly facilitates the biological interpretation among multiple integrated data sets. There is no requirement to pre-filter features in a study or map features from different data sets to a set of common genes. Therefore, MOGSA can be used to compare technology platforms that have different or missing features.

There is great potential for applying multi-table unsupervised GSA approaches for discovery of new subtypes and pathways in integrated data analysis of complex diseases such as cancer. In this study, we applied MOGSA in combination with clustering analysis. Dimension reduction approaches such as MOGSA and MFA are well suited to cluster discovery data because these approaches consider the global variance in the data and as such are complementary to hierarchical or k-means clustering approaches which focus on the pair-wise distance between observations (52–55).

The number of components is an important input parameter to consider when applying MOGSA to gene-set analysis or cluster discovery. Similar to PCA, the optimal number of MFA components may be assessed by examining the variance associated with each component. The first component will capture most variance and the variance associated with subsequent component decreases monotonically. Scree plots (Figure 2C, 4A) may be used to visualize if there is an elbow point in the eigenvalues, allowing one to select the components before the elbow point. Alternatively, one may select the number of components that capture a certain proportion of variance (50%, 70%, etc.). In addition, one may include components that are of biological interest. For example, in the iPS ES example, there is a clear biological meaning in the third component (ES vs iPS cell line). In the analysis of the BLCA data, we examined a range of components (1-12), and showed that there is little gain of information once a minimum number of components with high variance are included (Figure 6B). Users should also consider that the variance of retained components should not be dominated by one or a few data sets. To facilitate biological interpretation of components, the GSS could be decomposed with regard to components. In the BLCA example, the second and forth component are largely contributed by CNV, whereas mRNA is more important in defining the third and fifth components. Including five components ensured that both data sets contributed relatively similar variance to the total gene set score.

An issue might arise with latent variables analysis if components with large variance capture information unrelated to biological variance (51) such as technical artifacts or batch effects. In practice, this is rare in MFA, because it focuses on components that capture global correlation among all data sets. Often batch effects are specific to a platform and thus a component that captures information that is entirely uncorrelated to the global structure will be omitted from the set of highly variant integrated components. Despite that the decomposition of gene sets by components makes it is easy to remove unwanted variance, gene set scores calculated from the components reflecting variance of interest may lead to the better interpretation of interesting biology. However, it is still wise to perform careful batch effect control, especially in large scale omics studies. With moGSA, one may select non-contiguous components to enrich for specific patterns, or exclude attributes. For example we excluded the first component in the NCI60 cell line data which was associated with cell doubling time. One can perform linear regression analysis on the PCs (components) to generate GSS on features that are associated with a phenotype or clinical variable (eg stage, survival) of interest. A more detailed description of batch effect detection is described in (40). Another consideration when applying MOGSA is that it is most efficient in detecting gene-sets that have broad correlation patterns among data types. It may fail to discover gene-sets with few genes, particularly if they had low variances on the selected components.

Finally, MOGSA is computationally efficient, and performed comparable to ssGSA approaches ssGSEA and ssGSVA even when applied to large ‘omics integrative analysis of mRNA and CNV data of over 10,000 tumors. Open source, well documented code is provided in the Bioconductor package mogsa which includes a detailed vignette tutorial and example datasets. Whilst we do not include examples applied to multi omics single cell sequencing technologies that provide simultaneous measurement of transcriptomes and protein markers expressed in the same cell, (eg CITE-seq or REAP-seq (4, 5), MOGSA can be applied to integrating, interpreting and generating biological hypothesis from complex high dimensional data sets from bulk sequencing or single cell technologies.

## Acknowledgements

We wish to thank Prof. Joaquim Bellmunt for the insightful discussions about bladder cancer molecular subtypes and treatment. We also thank Hannes Hanne and Dominic Helm for reading the manuscript and giving valuable suggestions. Funding for this work was provided by DFCI BCB Research Scientist Developmental Funds, National Cancer Institute at the National Institutes of Health [grant numbers 2P50 CA101942-11, 1U19 AI111224-01, 1U19 AI109755-01] and Department of Defense BCRP [award number W81XWH-15-1-0013]. Views and opinions of, and endorsements by the author(s) do not reflect those of the US Army or the Department of Defense

## Description of additional data files

SupplementaryInformation.pdf – Supplementary methods and 23 supplementary Figures

Table_S1.xlsx - Table S1 - the gene-set score (GSS) matrix of Gene ontology (GO) for iPS ES 4-plex data.

Table_S2.xlsx - Table S2: The Chi square test of association between integrative subtypes and previously published subtypes.

Table_S3.xlsx - Table S3 - the gene-set score (GSS) matrix of Gene ontology (GO) and transcriptional factor target (TFT) gene-set with more than 200 significant GSSs for BLCA data.

Table_S4.xlsx - Table S4 - the gene influential score (GIS) for selected gene-sets (from Gene Ontology). The document contains GIS analysis for 9 gene-sets.

Table_S5.xlsx - Table S5 - the gene influential score (GIS) for selected transcriptional factor gene-sets. The document contains GIS analysis for 2 gene-sets.

## Abbreviations

ANOVA: analysis of variance;
AUC: area under the receiver operating characteristic curve;
BLCA: bladder cancer;
BP: biological process;
CC: cellular component;
CCA: canonical correlation analysis;
CIA: co-inertia analysis;
CLT: central limited theorem;
CPTAC: Clinical Proteomic Tumor Analysis Consortium;
DE: differentially expressed;
DEGS: differentially expressed gene-set;
EMT: Epithelial to mesenchymal transition;
GIS: gene influential score;
GO: gene ontology;
GS: gene-set;
GSA: gene-set analysis;
GSEA: gene-set enrichment analysis;
GSS: gene-set score;
MAD: median absolute deviation;
MCIA: multiple co-inertia analysis;
MF: molecular function;
MF: matrix factorization;
MFA: multiple factorial analysis;
MS: mass spectrometry;
MVA: multivariate analysis;
NMM: naïve matrix multiplication;
PCA: principal component analysis;
ROC: Receiver operating characteristic;
SVD: singular value decomposition;
TCGA: the cancer genome atlas;
TF: transcriptional factor;
TFT: transcriptional factor target

