## supplementary figures for "MOGSA: integrative single sample gene-set analysis of multiple omics data"

[$] Current address:

### Supplementary Methods

#### MOGSA algorithm

***Input data and gene-set annotation matrix***

The inputs to MOGSA are pairs of multiple matrices ($\mathbf{X}_K$, $\mathbf{G}_K$). $\mathbf{X}_K$, denotes a set of *K* matrices $\mathbf{X}_1$, ..., $\mathbf{X}_k$, … $\mathbf{X}_K$, are input matrices of omics data, where rows are features (e.g. genes, proteins) and columns are observations (e.g. cell lines, disease tissue samples). Matrix $\mathbf{X}_k$ have $p_k$ rows and all omics data have the sample number of $n$ columns. Each of the omics matrices $\mathbf{X}_1$, …, $\mathbf{X}_K$ has a corresponding gene-set annotation matrix, $\mathbf{G}_1$, ..., $\mathbf{G}_k$, …, $\mathbf{G}_K$. The gene-set annotation matrix $\mathbf{G}_k$ is a $p_k \times m$ binary incidence matrix of gene to gene-set membership associations, where $m$ is the number of gene-sets. The element $g_{k[i,j]}$ in $\mathbf{G}_k$ has the value 1 if the *i*th feature is a member of the gene-set *j* and 0 otherwise. $\mathbf{G}_k$ is constructed using predefined gene-set information such as the Gene Ontology [1, 2], GeneSigDb [3] or MSigDB [4].

***MOGSA step 1 multivariate integration***

The first step of the MOGSA involves data integration with a multiple table MF method. In this study, we use multiple factorial analysis (MFA) because of its simplicity and computational efficiency. MFA can be viewed as a generalization of principal component analysis (PCA) for a multi-table problem [5]. We briefly describe MFA using the nomenclature used in [5].

When integrating multiple data matrices, one must decide if all datasets should have equal weight, or if some data are “more important”, for example those with higher quality, fewer features, higher variance, etc. Simple tensor decomposition approaches, or PCA on a concatenated matrix, give every dataset equal weight and results are often dominated by the matrix (or matrices) with the large variance or most features. To correct for this, MFA weights datasets by dividing each by their first eigenvalue. The weight of each matrix is expressed as

$$\alpha_k = \frac{1}{\lambda_{k,1}^2} \qquad (1)$$

Where $\lambda^2_{k,1}$ is the first singular value of data matrix $\mathbf{X}_k$. For convenience, the weights of matrices are stored in a diagonal matrix **A**, whose diagonal elements are

$$\text{diag}\{\mathbf{A}\} = [\text{diag}\{\mathbf{A}_1\}, \cdots, \text{diag}\{\mathbf{A}_k\}, \cdots, \text{diag}\{\mathbf{A}_K\}] = [\alpha_1 \mathbf{1}_1^{\mathrm{T}}, \cdots, \alpha_k \mathbf{1}_k^{\mathbf{T}}, \cdots, \alpha_K \mathbf{1}_K^{\mathbf{T}}] \quad (2)$$

The transpose of a matrix is denoted by superscript $^{\mathrm{T}}$. $\mathbf{1}_k^{\mathrm{T}}$ is a vector of 1 in the length of $p_k$. As a result, **A** is a $p \times p$ diagonal matrix, the diagonal elements of **A** representing the weight of features in $\mathbf{X}_1$, ..., $\mathbf{X}_k$. Similarly, the weight of each observation is an $n \times n$ diagonal matrix, **M**. In the present study, we use $m_{ii}=1/n$, namely, all observations are equally weighted.

We then transpose and concatenate all $\mathbf{X}_k$ to a complete $p \times n$ matrix ( $p = \sum_k p_k$ ):

$$\mathbf{X} = [\mathbf{X}_1^{\mathrm{T}} \mid \ldots \mid \mathbf{X}_k^{\mathrm{T}} \mid \ldots \mid \mathbf{X}_K^{\mathrm{T}}]^{\mathrm{T}} \quad (3)$$

After deriving the matrix weights, observation weights and the concatenated matrix, MFA is reduced to an analysis of the triplet (**X**, **A**, **M**). The solution of the problem is given by generalized singular value decomposition (GSVD):

$$\mathbf{X}^{\mathrm{T}} = \mathbf{P} \Delta \mathbf{Q}^{\mathrm{T}} \text{ with the constraint that } \mathbf{P}^{\mathrm{T}} \mathbf{M} \mathbf{P} = \mathbf{Q}^{\mathrm{T}} \mathbf{A} \mathbf{Q} = \mathbf{I} \quad (4)$$

**X** is transpose so that **P** is an $n \times r$ matrix, **Q** is a $p \times r$ matrix, $\mathbf{\Delta}$ is an $r \times r$ square matrix. The maximum number of $r$ is the rank of **X**. The matrix storing components of MFA, **F**, are given by

$$\mathbf{F} = \mathbf{P} \mathbf{\Delta} \quad (5)$$

where **F** has the same dimension as **P**. In the PCA framework, the matrix **P** contains the PCs or latent variables. We also call it *sample space* in this paper because every row in **P** corresponds to a sample in **X**. The matrix **Q** is the loading matrix or *feature space* as every row in P corresponds to a feature. Because **X** is a concatenation of multiple matrices, the feature space matrices **Q** is also a concatenation of multiple $\mathbf{Q}_k$ matrices, namely,

$$\mathbf{Q} = [\mathbf{Q}_1^{\mathrm{T}} \mid \cdots \mid \mathbf{Q}_k^{\mathrm{T}} \mid \cdots \mid \mathbf{Q}_K^{\mathbf{T}}]^{\mathbf{T}} \quad (6)$$

***MOGSA step 2 project gene-set annotation matrix as supplementary data***

Different gene-sets have different candidate genes, therefore, in order to facilitate the comparison of gene-set score across gene-sets, we normalized the gene-set annotation matrix so that the sum of each column in **G**, where $\mathbf{G} = [\mathbf{G_1}^T|\ldots|\mathbf{G_k}^T|\ldots|\mathbf{G_K}^T]^T$, equals 1, that is,

$$\hat{g}_{[i,j]} = \frac{g_{[i,j]}}{\sum_i g_{[i,j]}} \tag{7}$$

where $\hat{g}_{[i,j]}$ is the elements on the *i*th row and *j*th column in the normalized gene-set annotation matrix $\hat{\mathbf{G}}$ ($p\times m$), where $\hat{\mathbf{G}} = [\hat{\mathbf{G}}_1^T | \cdots | \hat{\mathbf{G}}_k^T | \cdots | \hat{\mathbf{G}}_K^T]^T$. The gene-set score can be calculated using either un-normalized or normalized gene-set annotation, but we will use the normalized version to describe the method.

Next, we project the annotation matrix as supplementary data to generate the *gene-set space* matrix $\mathbf{W}_k$ ($m\times r$) [6], which is calculated as a product of the normalized gene annotation matrix and loading matrix.

$$\mathbf{W} = \hat{\mathbf{G}}^T \mathbf{AQ} \tag{8}$$

The overall gene-set space **W** ($m\times r$ matrix) could also be expressed as the sum of individual $\widehat{\boldsymbol{W}}_k$, that is,

$$\mathbf{W} = \sum_{k=1}^{K} \mathbf{W}_k \text{ where } \mathbf{W}_k = \hat{\mathbf{G}}_k^T \mathbf{A}_k \mathbf{Q}_k \tag{9}$$

***MOGSA step 3 reconstruction of gene-set-observation matrix***

The main output of MOGSA is a *gene-set score (GSS)* matrix, denoted by **Y**, whose rows are *m* gene-sets and columns are *n* observations. It is calculated as

$$\mathbf{Y} = \hat{\mathbf{G}}^T \mathbf{AQ}^{[R]} \mathbf{\Delta}^{[R]} \mathbf{P}^{[R]T} = \mathbf{W}^{[R]} \mathbf{F}^{[R]T} = \hat{\mathbf{G}}^T \mathbf{AX}^{[R]} \tag{10}$$

where $\mathbf{Q}^{[R]}$ and $\mathbf{P}^{[R]}$ are the gene space and observation space within top *R* components. $\mathbf{\Delta}^{[R]}$ is the diagonal matrix containing top *R* singular values. As a result, $\mathbf{X}^{[R]}$ is the reconstruction of **X** using top *R* components.

In practice, it is interesting to know which dataset or component contribute more to the overall gene-set score. Therefore, we decompose gene-set scores with respect to data sets and components. The GSS matrix for dataset $\mathbf{X}_k$ and component $r$ is calculated as

$$\mathbf{Y}_k^r = \mathbf{W}_k^r \mathbf{F}_k^{r\,\mathrm{T}} \tag{11}$$

we use superscript $r$ to indicate the $r$th component and the subscript $k$ to indicate the $k$th matrix ($\mathbf{X}_k$). Similarly, $\mathbf{W}_k^r$ denotes the $r$th dimension of gene-set space of matrix $\mathbf{X}_k$, $\mathbf{F}_k^r$ is the $r$th component of the sample space. The outer product of the two vectors results in a GSS matrix for a specific components and dataset. Consequently, the overall gene-set score for component $r$ (i.e. component-wise decomposed gene-set scores) is the sum of the gene-set score matrix of the components across all datasets, that is,

$$\boldsymbol{Y}^r = \sum_k \boldsymbol{Y}_k^r = \sum_{k=1}^{K} \boldsymbol{W}_k^r \boldsymbol{F}_k^{r\,\mathrm{T}} \tag{12}$$

Similarly, the overall gene-set score matrix by a single dataset (i.e. data-wise decomposed gene-set scores) is the sum of the matrices by all the components retained.

$$\boldsymbol{Y}_k = \sum_r \boldsymbol{Y}_k^r = \sum_{r=i}^{R} \boldsymbol{W}_k^r \boldsymbol{F}_k^{r\,\mathrm{T}} \tag{13}$$

Therefore, the contribution of an individual dataset and/or component may be calculated. Finally, the complete gene-set score matrix is given by

$$\mathbf{Y} = \sum_r \mathbf{Y}^r = \sum_k \mathbf{Y}_k = \sum_{k=1}^{K} \sum_{r=1}^{R} \mathbf{W}_k^r \mathbf{F}_k^{r\,\mathrm{T}} \tag{14}$$

which is the sum of all contributions by individual components and dataset. In practice, only the components with greatest variances (highest eigenvalues) should be retained in the analysis. If all components are retained, the result would be similar or exactly the same as naïve matrix multiplication (NMM; see later).

*Evaluation of the significance of gene-set scores (calculating p-values)*

The expression (7) and (10) say that, for each observation, a gene-set score could be viewed as the weighted mean of gene expression (in the reconstructed expression values $\mathbf{X}^{[R]}$) of genes in a particular gene-set.

If the candidate genes in a gene-set are randomly drawn from all features in $\mathbf{X}^{[R]}$ (null hypothesis), the distribution of the means of selected genes is given by central limited theorem (CLT),

$$\bar{x} \sim N(\mu, \sigma_{\bar{x}}) \text{ with } \sigma_{\bar{x}} = c\frac{\sigma}{\sqrt{h}} \tag{15}$$

Where $\mu$ is the mean of a column (observation) in $\mathbf{X}^{[R]}$, $\sigma_{\bar{x}}$ is the sampling standard deviation of means, $\sigma$ is the standard deviation of the column in $\mathbf{X}^{[R]}$, $h$ is the number of candidate genes mapped to **X** in a gene-set and $c = \sqrt{(p-h)/(p-h)}$ is the finite population correction factor ($p$ is the number of features in **X**). The finite population correction factor is used as each gene was only selected once in one gene-set.

***Gene influential score***

Gene-sets are composed of genes, therefore, it is also important to evaluate the contribution of each feature to the GSS. The genes with large contribution could be view as "driver" genes in a gene-set. In MOGSA, feature contribution, denoted by gene influential score (GIS), is calculated via a leave-one-out procedure.

The GSS of gene-set $i$, $\mathbf{Y}_{[i]}$, for all the observations are

$$\mathbf{Y}_{[i]} = \hat{\mathbf{G}}_{[i]}^{\mathrm{T}}\mathbf{A}\mathbf{X}^{[R]} \tag{16}$$

where $\hat{\mathbf{G}}_{[i]}$ is the gene-set annotation vector for gene-set $i$. Correspondingly, the gene-set score for $i$th gene-set excluding gene $g$ is

$$\mathbf{Y}_{[i]}^{-g} = \hat{\mathbf{G}}_{[i]}^{-g\,\mathrm{T}}\mathbf{A}\mathbf{X}^{[R]} \tag{17}$$

Where $\hat{\mathbf{G}}_{[i]}^{-g}$ is the gene-set annotation vector for gene-set $i$ but without gene $g$. The influence of the gene $g$ is measured by

$$E_{[i]}^{g} = -\log_{2} \frac{sd(\mathbf{Y}_{[i]}^{-g})}{sd(\mathbf{Y}_{[i]})} \tag{18}$$

where $sd(\cdot)$ stands for the function of calculating standard deviation. For convenience, the feature influential score then is rescaled, such that the gene with maximum influence always equals 1. In general, a positive $E_{[i]}^{g}$ suggests that gene *g* tends to have a positive correlation with gene-set score of gene-set [*i*], whereas a gene with a negative value tends to have a negative correlation.

**Other GSA methods and Naïve Matrix multiplication (NMM)**

Single gene-set method, including GSVA and ssGSEA methods were implemented using the R/Bioconductor package GSVA [7]. Default settings were used for these methods. Naïve gene-set score $\mathbf{Y}_{\mathbf{naive}}$ was calculated through matrix multiplication (NMM).

$$\mathbf{Y}_{naive} = \hat{\mathbf{G}}^{\mathrm{T}}\mathbf{X} \tag{21}$$

Therefore, the result of NMM is exactly the same as MOGSA if all of the axes are retained.

### Supplementary figures

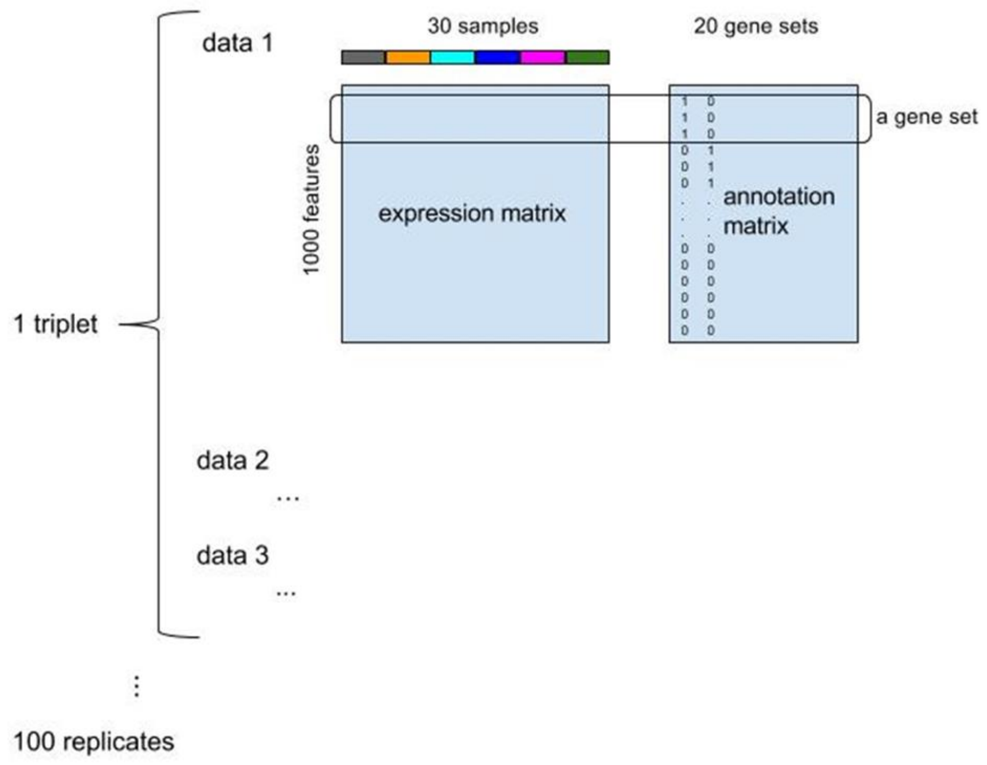

**Figure S1 – A diagram showing the data simulation data.**
One dataset contains a matrix triplet (data 1, data 2 and data 3). Each triplet contains 1,000 features and 30 observations. The 30 observations were divided into six clusters, 5 observations in each cluster. The 1,000 features are assigned to 20 gene-sets (each gene-set had 50 genes), coded in the gene set annotation matrix. 100 triplets were simulated in this analysis.

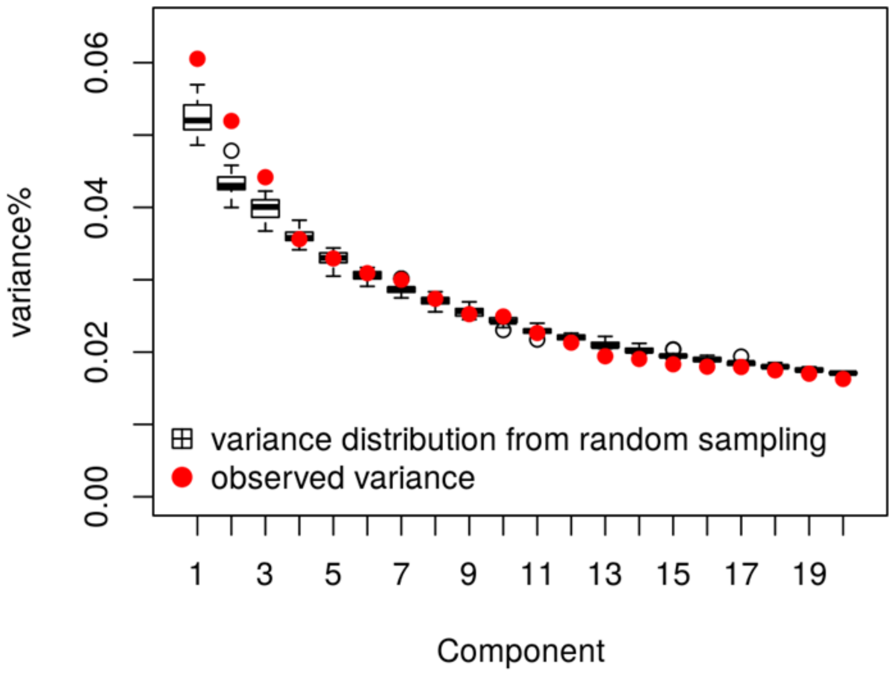

**Figure S2 – Determining the number of components that capture the correlated structure between NCI60 transcriptome and proteome data**

A random sampling method was used to determine the number of components capturing significant correlated structure and between transcriptome and proteome. To this end, the cell lines labels were randomly shuffled in both transcriptomic and proteomic data and the variance of components were calculated from the randomly labels data. We preformed this process 20 times in order to estimate the null distribution of variances associated with each component and found that the variance of top three components are significantly higher than the null distribution. In **Figure S12,** we show that component 1 was significantly correlated with cell doubling time.

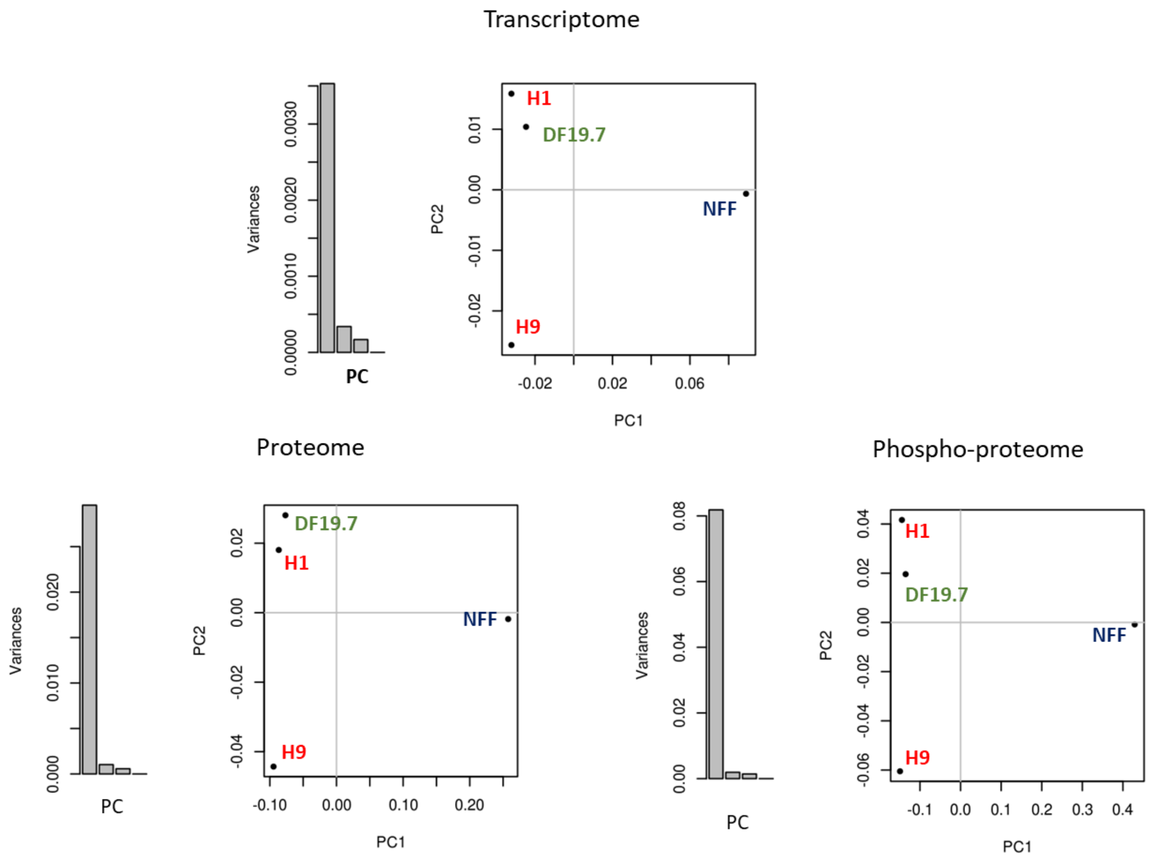

**Figure S3 – PCA of transcriptomic, proteomic and phosphoproteomic data of iPS/ES cell lines**
Most of the variances are captured by the first component which reflect the difference between the fibroblast foreskin cells (NFF) and the other cell lines.

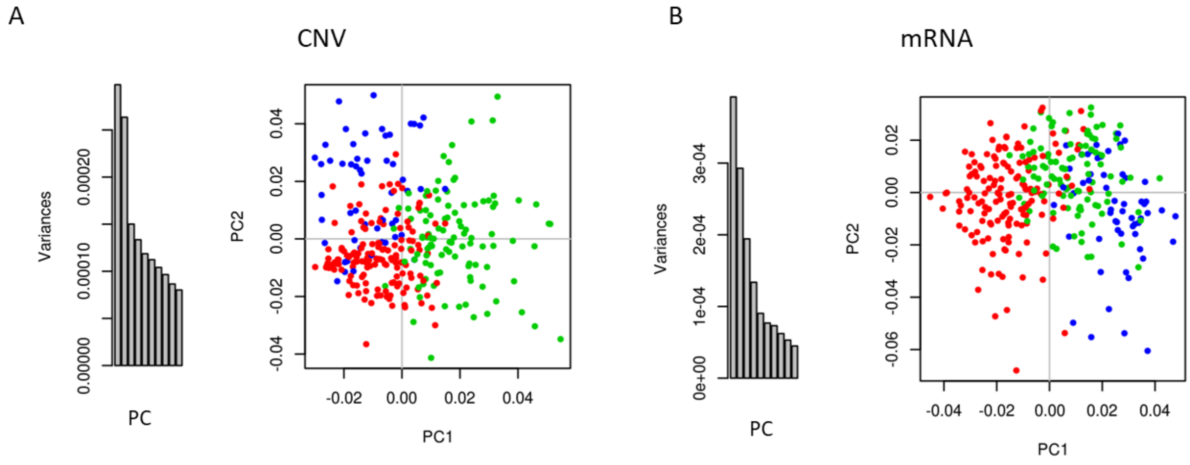

**Figure S4 – PCA of (A) CNV and (B) transcriptomic data of BLCA tumors (n=308)**
Each panel shows a scree plot of the variance captured by the first 10 components and a plot of the first two components (PC1, PC2). Tumors are colored by molecular subtype; C1 (red), C2 (green) and C3 (blue). The first two components of the CNV decomposition distinguishes these 3 subtypes. The first eigenvalue (square of singular value) of transcriptomic and CNV data were 0.0004 and 0.0003 respectively, which values were used to calculate the weight of each dataset in MFA.

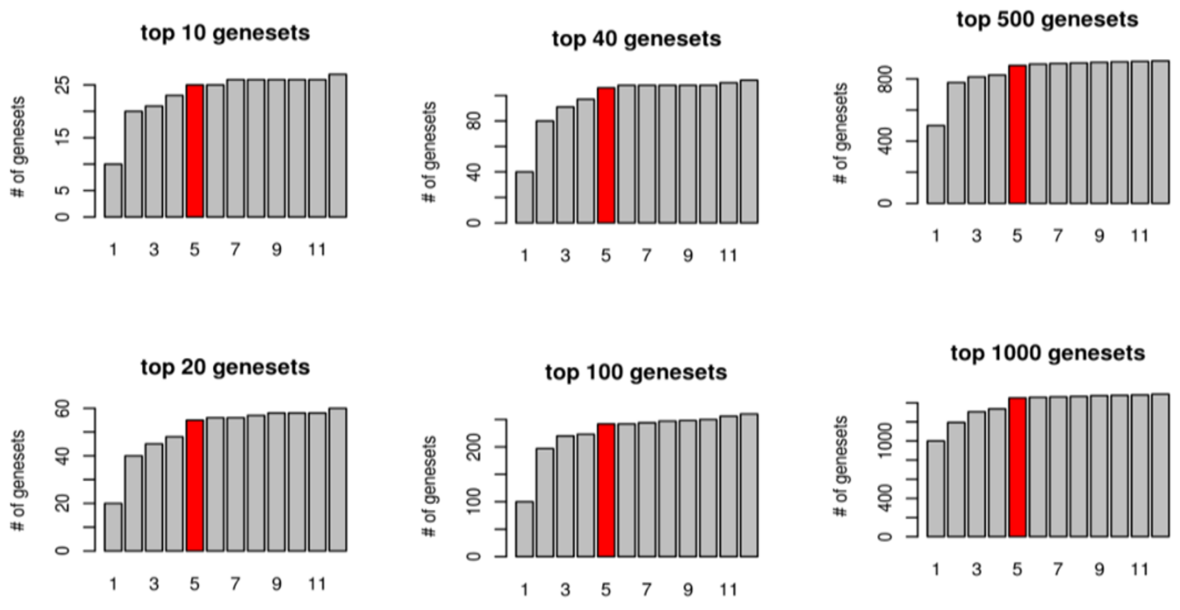

**Figure S5 – Five components identifies robust top-ranking gene sets**

Gene set scores (GSS) were calculated when 1 to 12 components were included. In every cases (1-12 components), gene sets were ranked from high to low according to the number of patients in which their GSSs was significant. Next, top N (N=10, 20, 40, 100, 500, 1000) gene sets were selected. The figure shows how the union of gene sets increases when additional components are examined. Using the top left panel as an example, with one component we extracted 10 gene sets, when we add a second components in the calculation of GSSs, we extracted another 10 gene sets which are completely different with ones identified using a single component, results in total 20 gene sets. In general, the figure shows that 5 component is an elbow point at which number including more component does not result in distinct top-ranking gene sets.

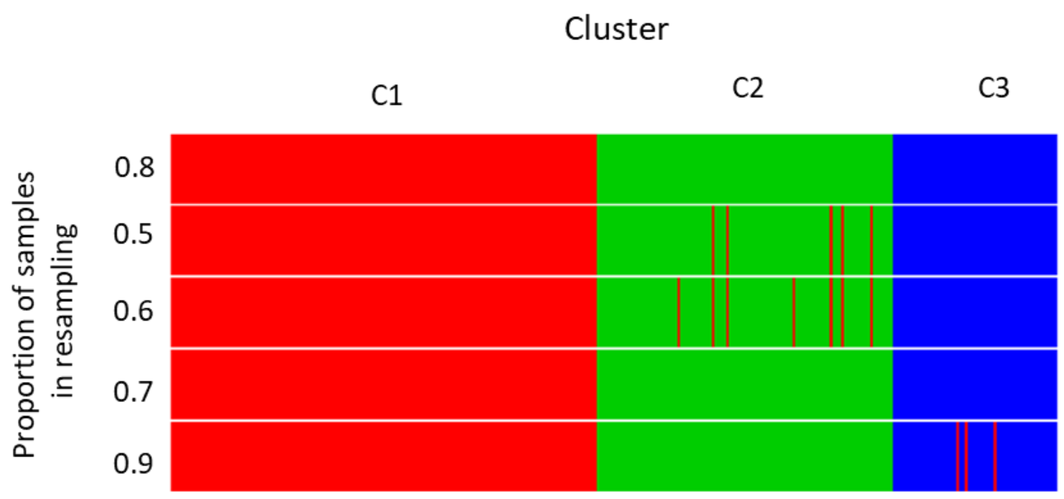

**Figure S6 - Stability of consensus clustering result when different resampling size is used** where C1 (red) C2 (green) C3 (blue) are indicated by color bars.

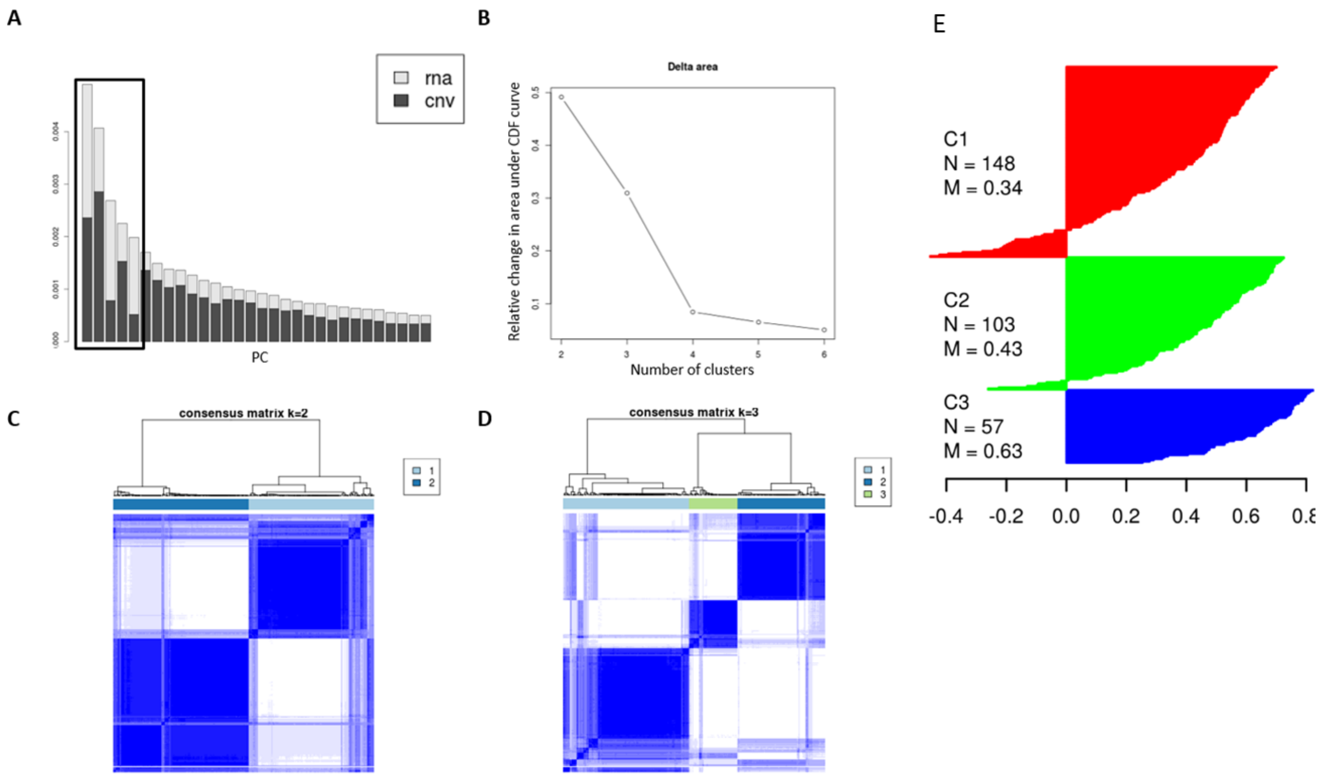

**Figure S7 – Clustering of MFA latent variables identify three BLCA subtypes**
MFA of mRNA and CNV of BLCA patients was performed. (A) shows the eigenvalues of each of the latent variables and top five PCs are marked. Five latent variables were used in consensus clustering and (B) the relative change area under the CDF curve (y-axis) over different pre-defined number of clusters (x-axis), which is used to determine the number of clusters. For both 2 and 3 clusters, the relative change in area under the CDF cure is high, indicating that the BLCA tumors may contain either 2 or 3 subtypes. Hierarchical of the consensus matrix for (C) 2 or (D) 3 subtypes. (E) silhouette analysis of three clusters.

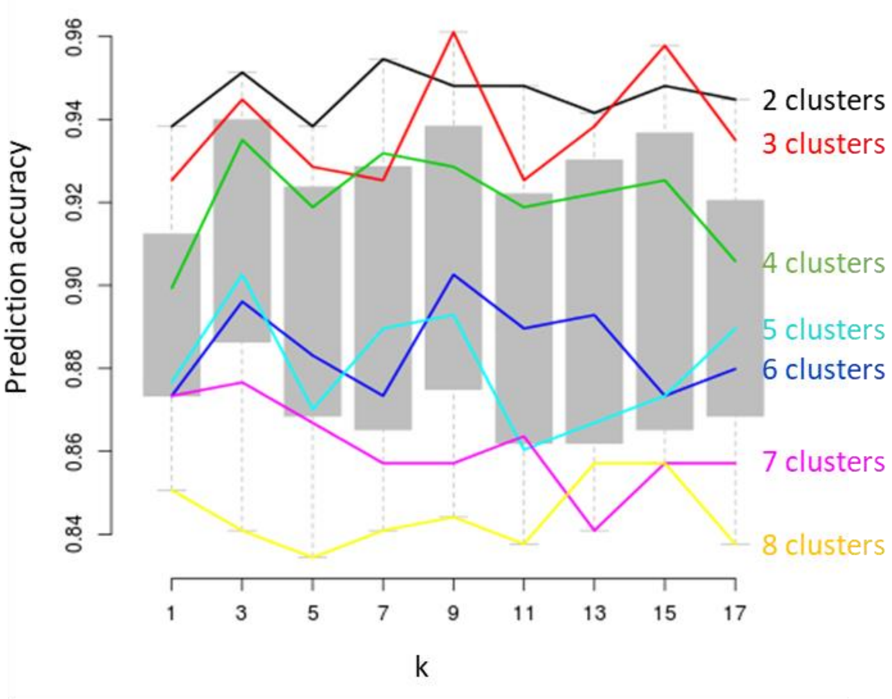

**Figure S8 – Determining the number of K in KNN algorithm (Use in calculating prediction Strength)**

Cross-validation were used to optimize the optimal number of K in the KNN classifier. We evaluated odd numbers K from 1 to 17. The performance of classifier were measured with prediction accuracy (y-axis). There is not a K clearly better than the others.

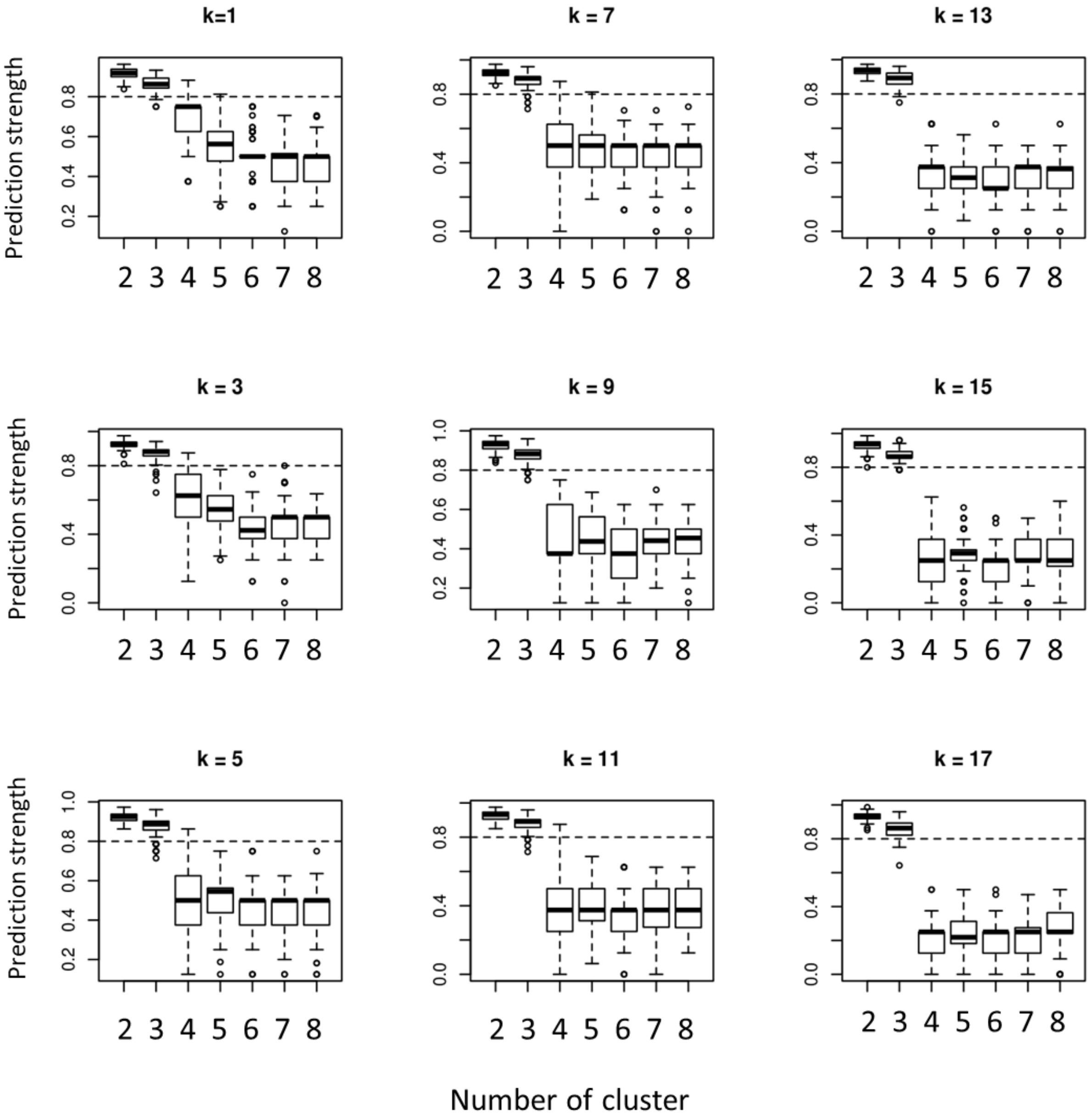

**Figure S9 – Prediction strength using different K in KNN classifier.**
All K suggest that three subtype is the robust number of subtype in the integrated BLCA datasets.

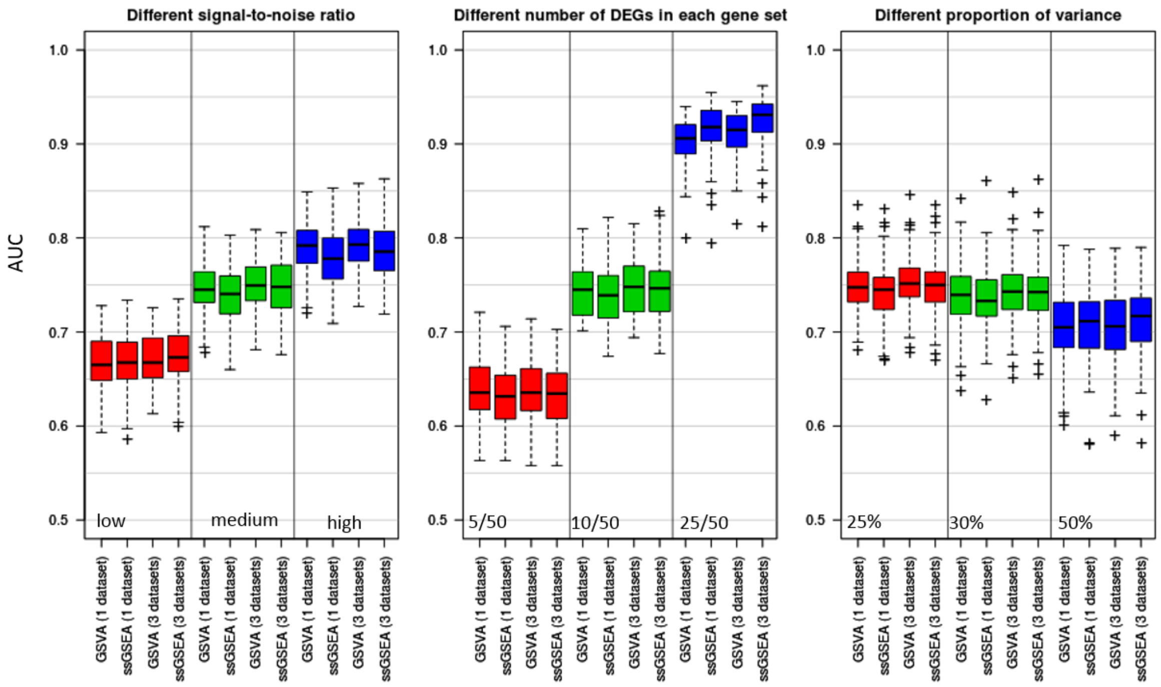

**Figure S10 – Simple concatenation of multiple data sets did not improve the performance of GSVA and ssGSEA**

The plots show area under the curve (AUC) of performance of GSVA and ssGSEA analysis of a single dataset (referred as 1 data set) and concatenated data sets (referred as 3 data sets). Methods, data and evaluation are the same those in Figure 2.

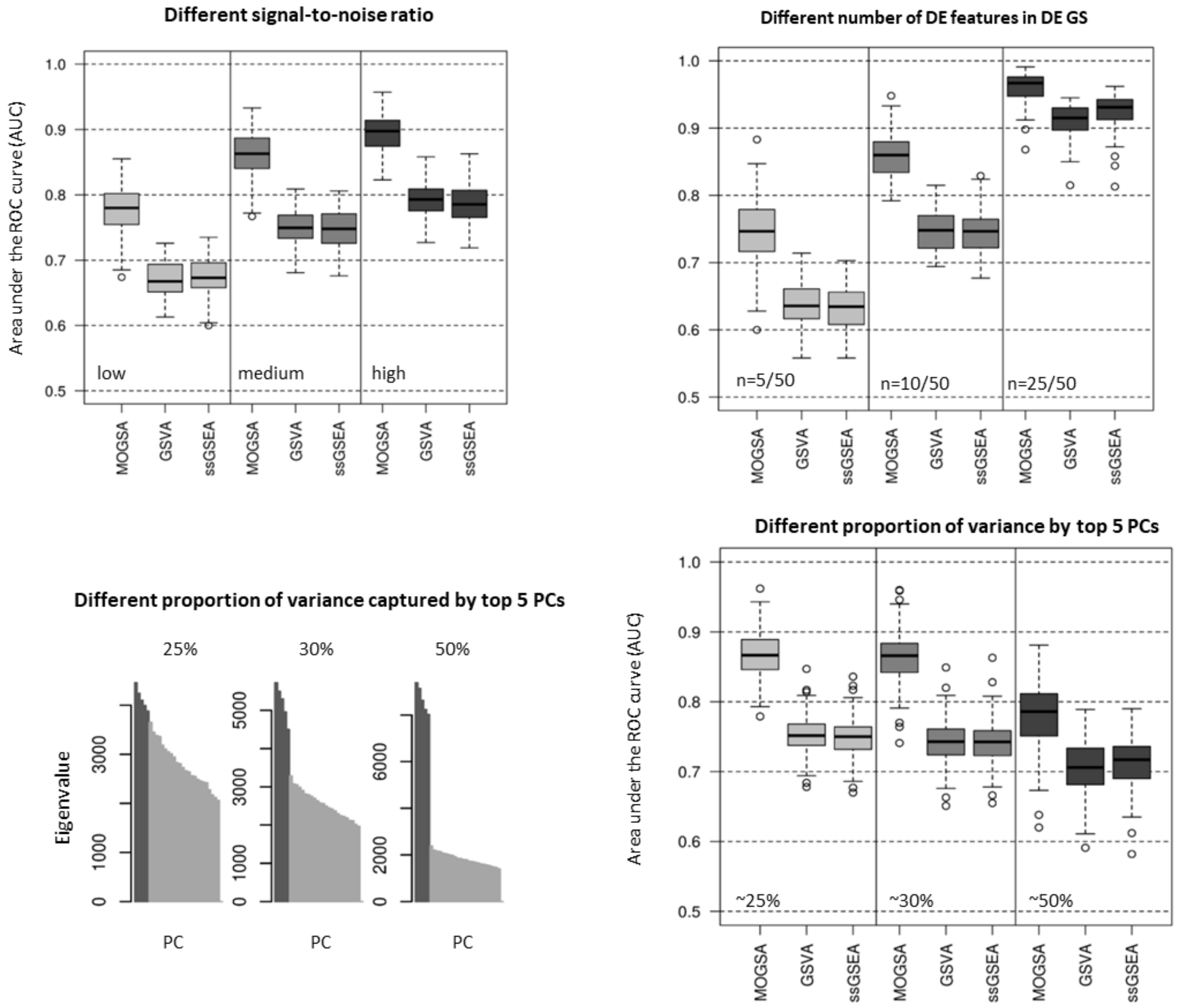

**Figure S11 – MOGSA outperforms GSVA and ssGSEA using weighted matrices**

Because MOGSA weights input matrices according to their first singular value, we weighted the matrices in a triplet by their first singular value before concatenation, MOGSA still outperforms others. The analyses are the same as described in Figure 2.

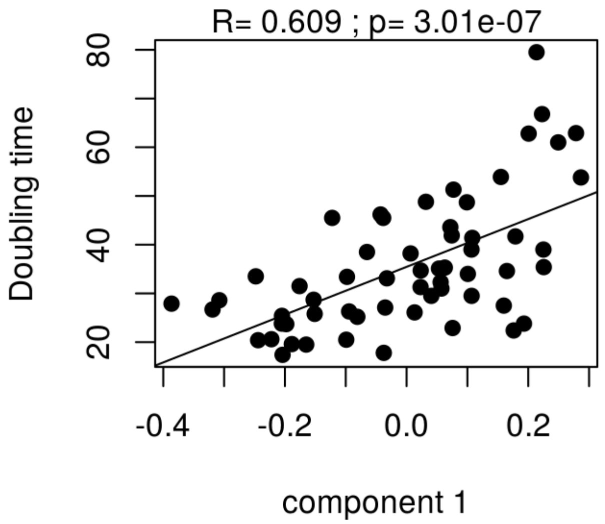

**Figure S12 – Component 1 of NCI60 data is significantly correlated with cell doubling time**

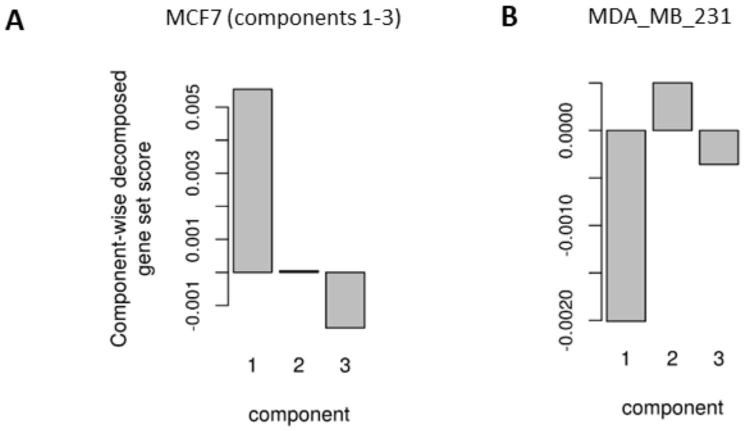

**Figure S13 – the component-wise decomposed gene set score of cycle checkpoint pathway in MCF7 and MDA_MB_231 cell line.**

Component 1 is the driving force of significant level in these two cell lines.

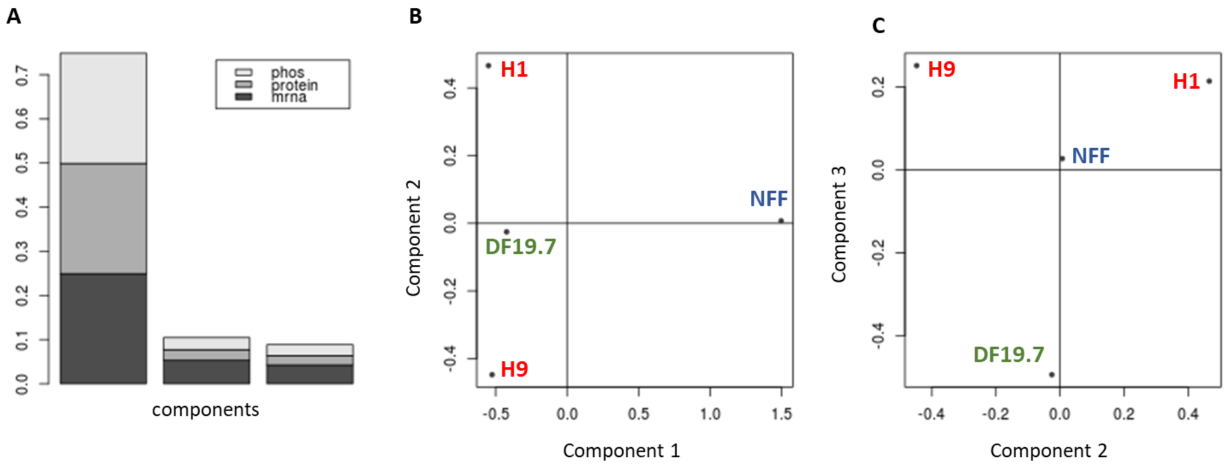

**Figure S14 – MOGSA of the iPS ES 4-plex data**
(A) A scree plot of the eigenvalues of the MFA. Grayscale shades represent the contribution of each individual dataset. Each data set contributes roughly equally to the total variance of a component. Similar to PCA of the individual datasets, the first component captures most of the variance in the data. (B) shows that the first component captures the difference between NFF and pluripotent cell lines and (C) shows the third component represents the difference between iPSC (DF19.7) and ESC lines.

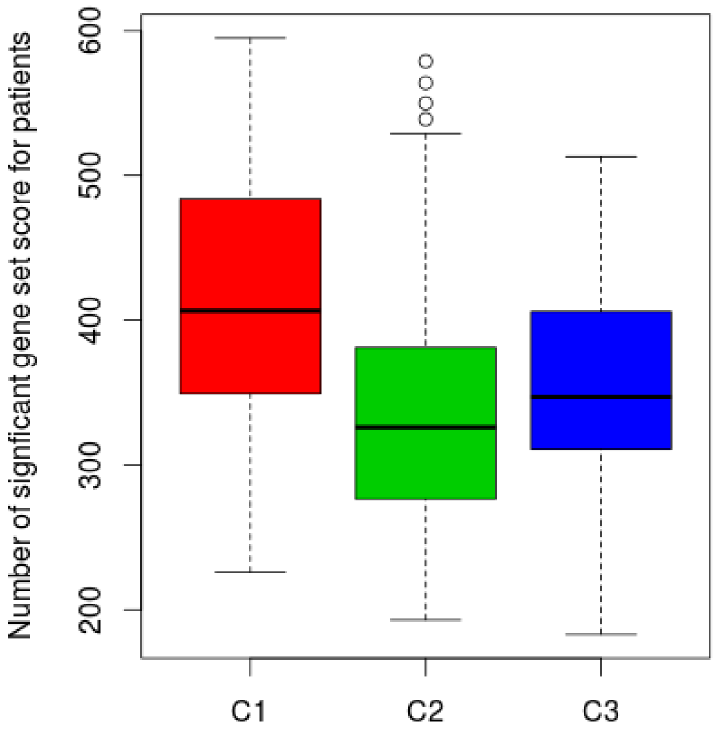

**Figure S15 - The number of significant gene set scores (GSS) per patient**
Number of genesets with either positive or negative significant GSS scores in BLCA subtypes

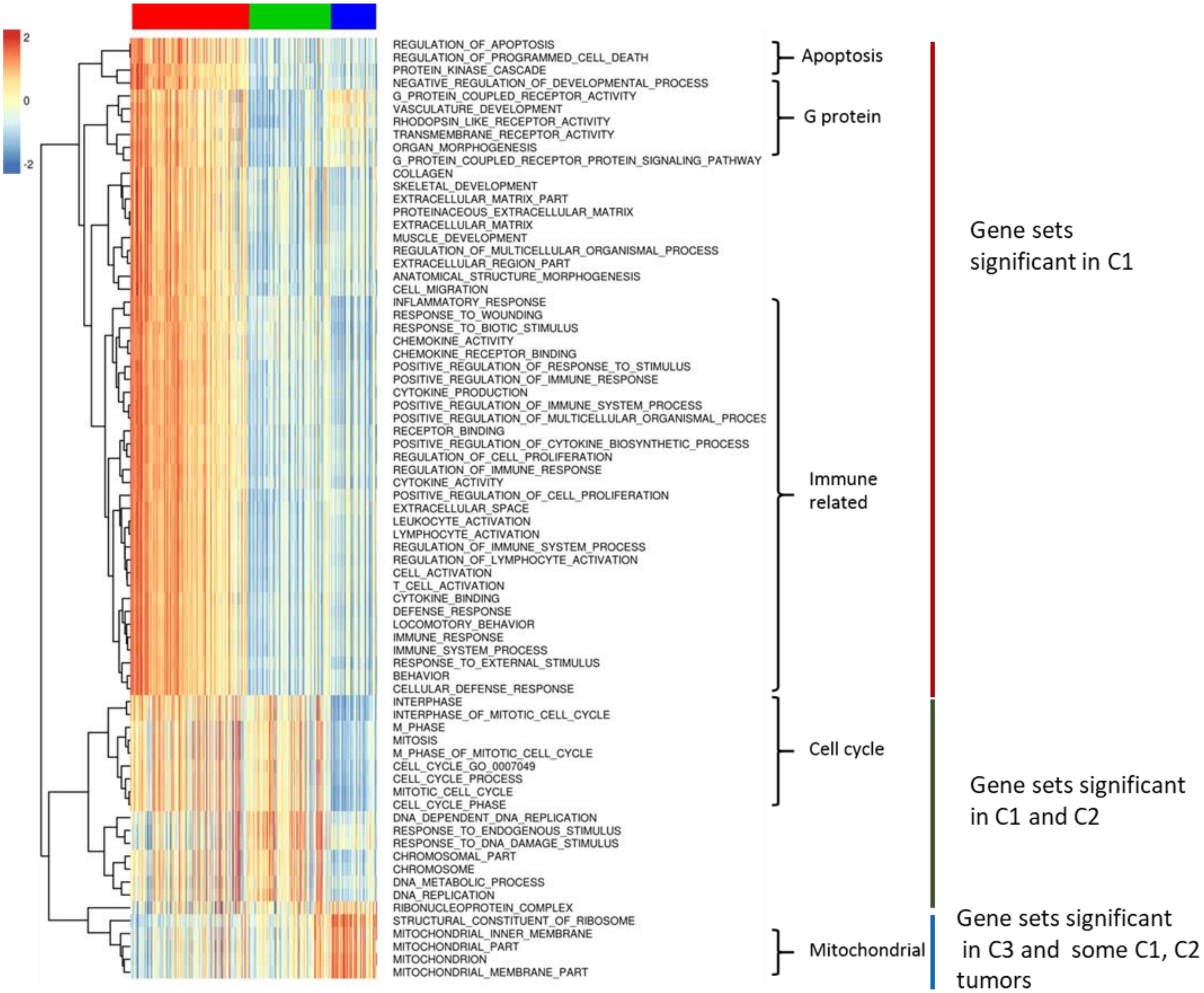

**Figure S16 – Gene sets that are significantly up or down regulated in more than 200 patients in BLCA**

The rows (gene sets) of the heatmaps are clustered so that the gene sets with similar GSS scores across patients are grouped. Columns are ordered according to BLCA tumor molecular subtype (C1, C2 and C3). Gene sets formed three broad clusters (those significant in C1, C1 and C2 or C3 and other tumors). Significant gene sets in C1 were associated with apoptosis, G protein coupled proteins, extracellular function, muscle development and Immune response. Gene sets significant in both C1 and C2 were mostly associated with the cell cycle, DNA repair and replication. Gene sets significant in C3 patients were associated with the mitochondria.

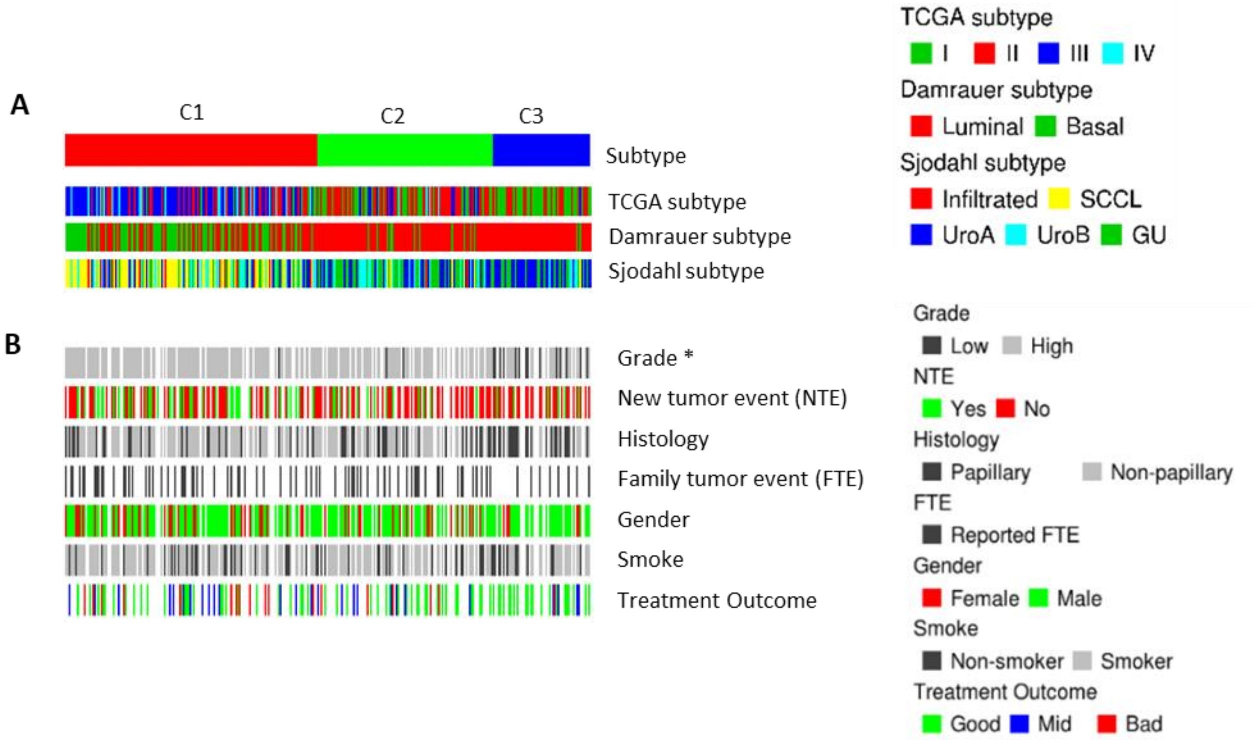

**Figure S17 – Characteristics of the BLCA molecular subtypes.**
(A) There was strong concordance between the integrative subtypes and molecular subtypes previously reported by the TCGA, Damrauer et al. and Sjodhl et al. C1 was enriched with III and IV for TCGA subtype, Basal subtype in Damrauer subtype and the SCCL and Infiltrated subtypes in Sjodahl subtype. C2 and C3 is comparable to the luminal subtype in Damrauer subtype model. C3 also enriched with UroA subtype in Sjodahl subtype and type I in TCGA subtype model. (B) Enrichment of clinical/phenotype factors including smoking gender, new tumor events, etc ib subtypes was studies. Grade was significantly correlated with the subtypes ($\chi2$ test, FDR BH corrected p value $< 0.01$).

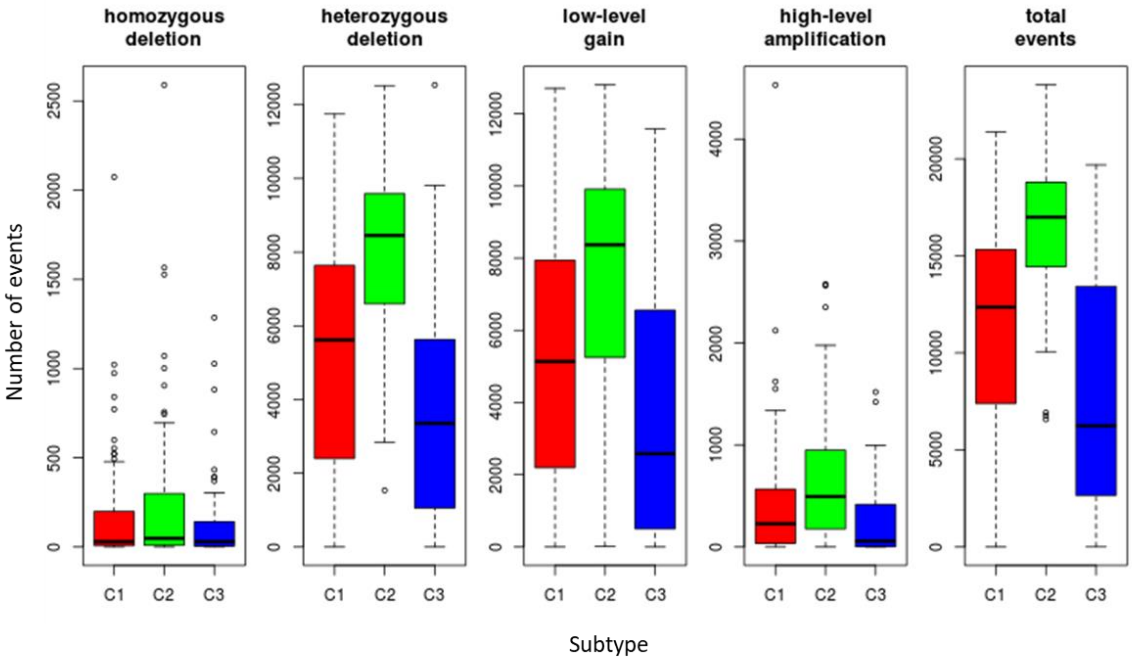

**Figure S18– BLCA subtype C2 has more instability and higher numbers of mutation events**
The figure shows the numbers of homozygous or heterozygous deletions, low and high level gains in addition to total CNV events in the genome of BLCA patients (n=308).

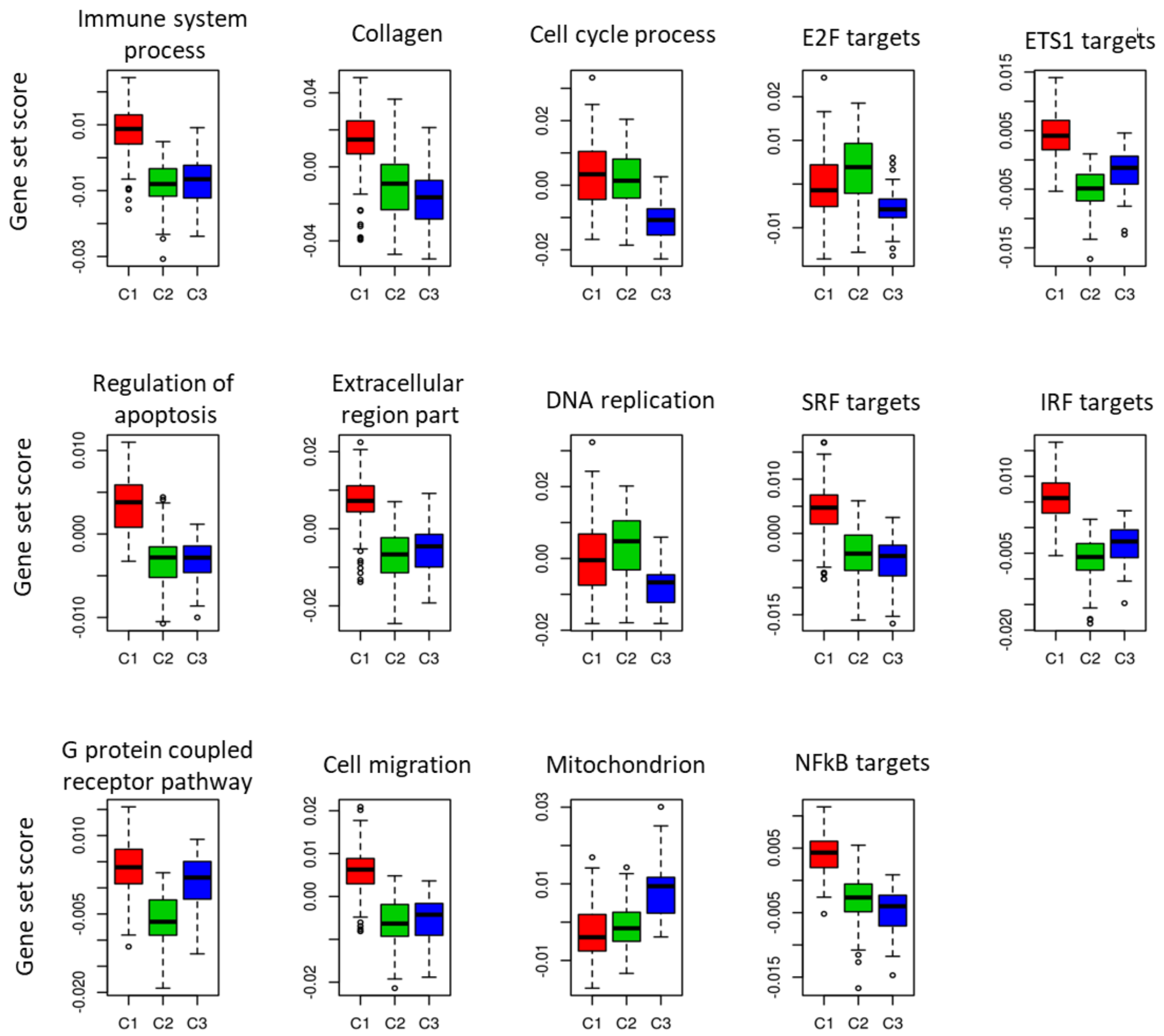

**Figure S19 – The distribution of gene set scores in different BLCA molecular subtypes**

Boxplot of gene sets scores in figure 5 C and D. Immune processes, Regulation of apoptosis, and cytosketal gene sets were upregulated in C1. C2 was characterized by downregulation of ETS1 and IRF targets, G-protein coupled receptors pathways and increased DNA related pathway (possibly associated with increased genome instability). C3 had lower expression of cell cycle and DNA replication genes compared to C1 and C2.

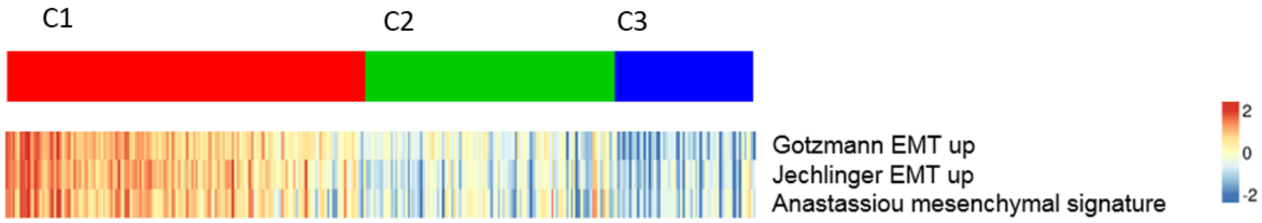

**Figure S20 –EMT related gene sets are highly activated in the C1 subtype.**
Heatmap displays GSS for three mesenchymal related gene sets downloaed from MSigDB C2 curated signatures. The original names as annotated in the MSigDB are: "GOTZMANN_EPITHELIAL_TO_MESENCHYMAL_TRANSITION_UP","JECHLINGER_EPITHELIAL_TO_MESENCHYMAL_TRANSITION_UP" ,"ANASTASSIOU_CANCER_MESENCHYMAL_TRANSITION_SIGNATURE".

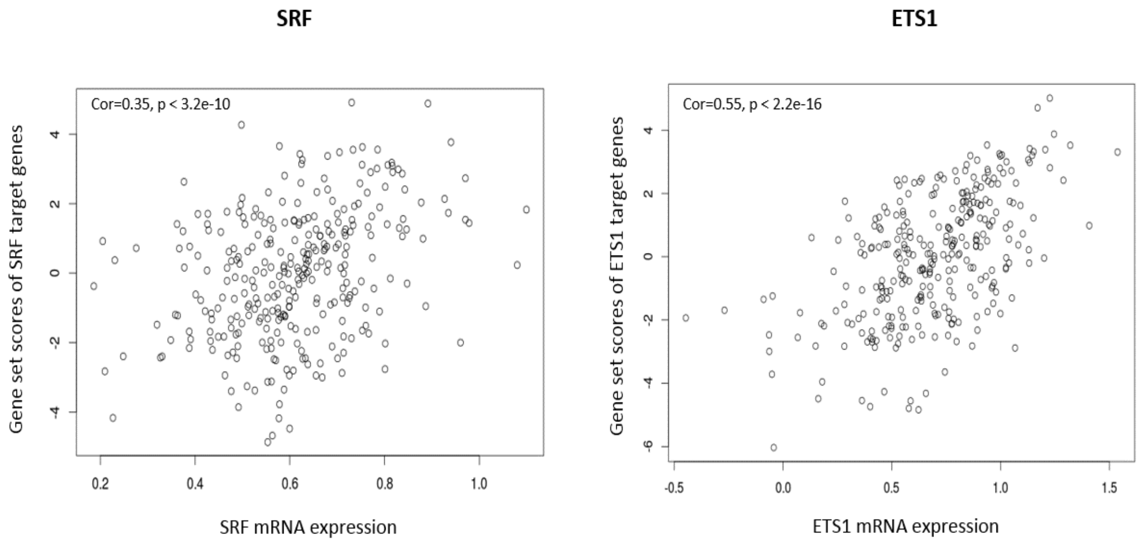

**Figure S21 – Gene sets scores of transcription factor (TF) target gene sets were highly correlated with the mRNA expression of their transcript factors in tumors**
Scatter plots show gene set score and mRNA expression levels of transcription factors (A) SRF and (B) ETS1 in the 308 BLCA tumors.

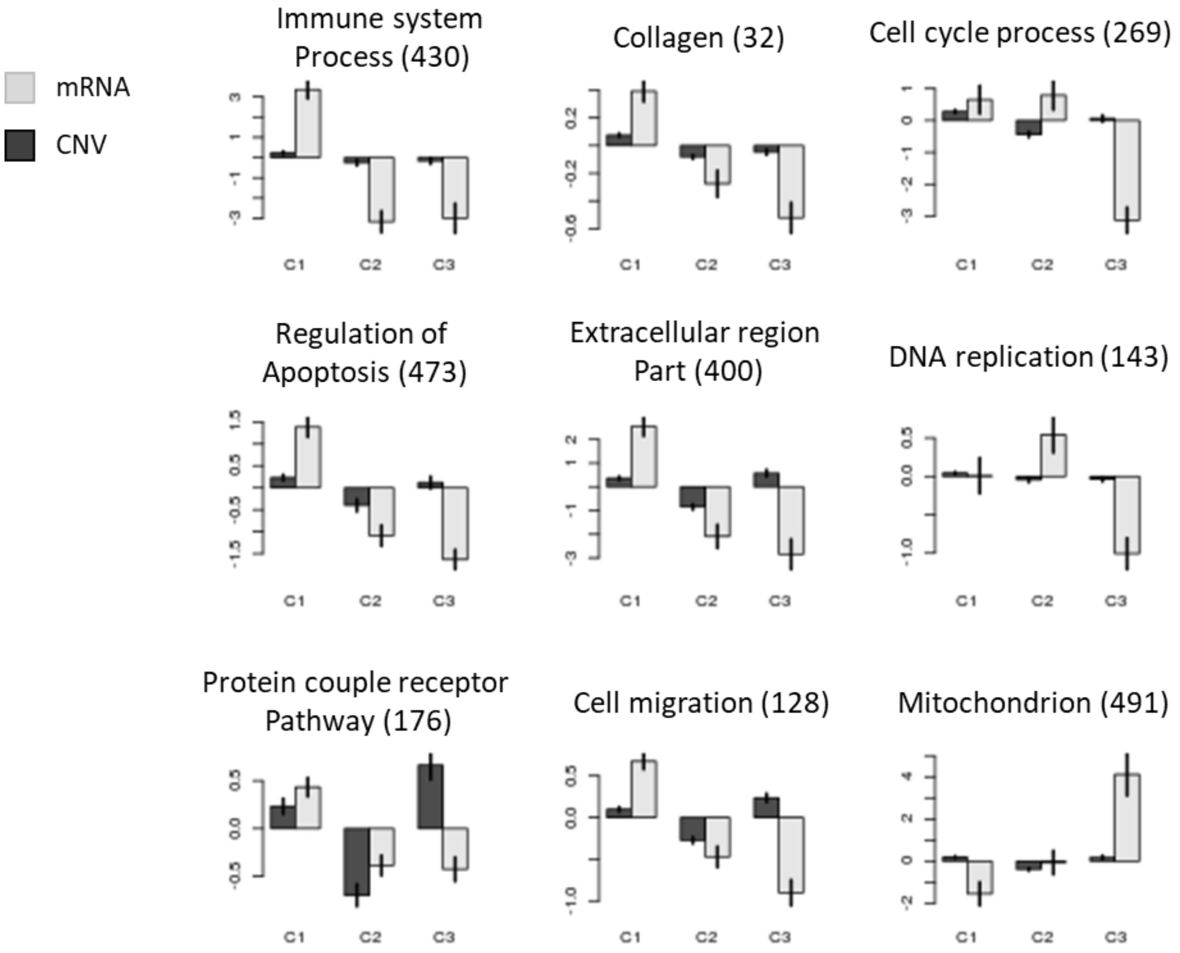

**Figure S22 – Non-normalized gene set scores of gene sets that were significant in BLCA tumors (related to Figure 5C)**

Plots are labelled with the gene set name and the number features (genes) in each gene set which is shown in parenthesis. The raw GSS is sum of the contributions of genes in a gene set. Therefore, the scale of non-normalized gene set scores (y-axis) are different. Gene sets with more genes will have higher scores so that GSS will not be comparable within a study. To solve this problem, MOGSA normalizes raw GSS by gene set size. Normalized GSS are reported throughout this article.

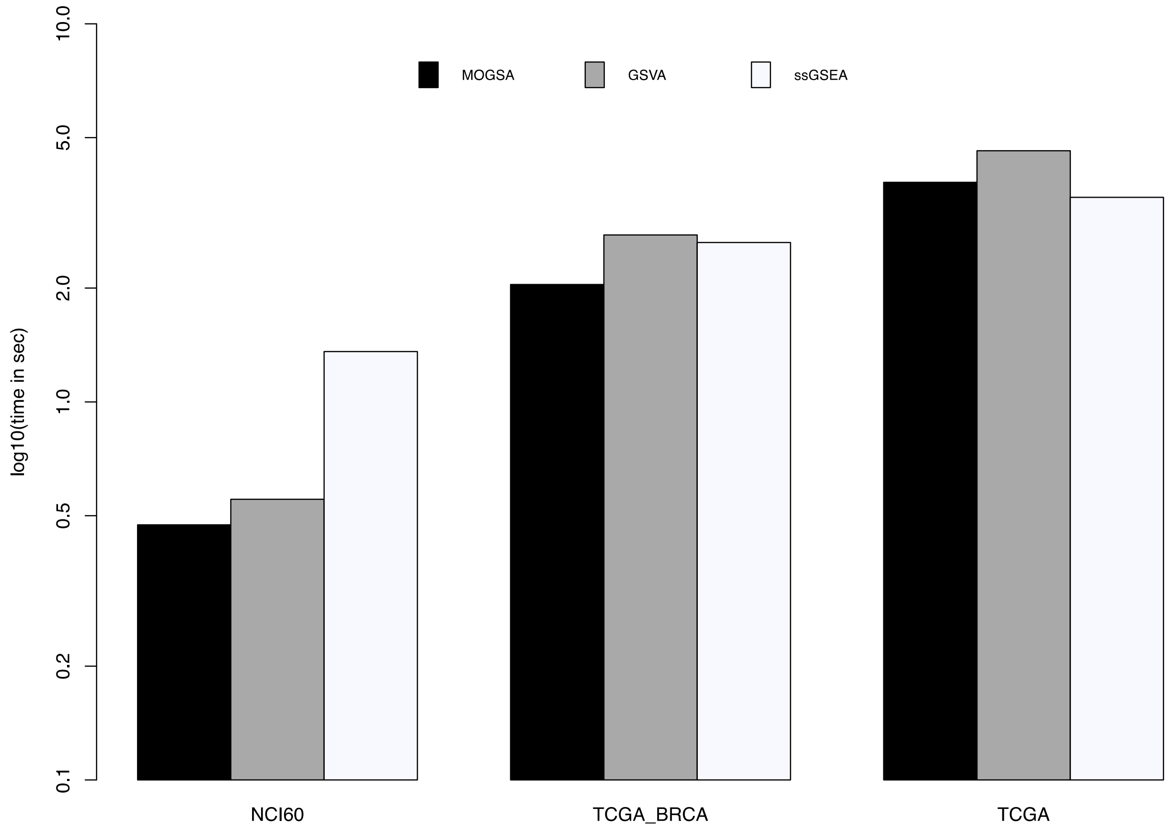

**Figure S23 – Computational efficiency of MOGSA**

Computational time to perform ssGSA using MOGSA, GSVA, ssGSEA on different size datat sets; NCI60 cell line transcriptomic datasets (58 cell lines, 17967 genes, 50 Hallmark genesets), mRNA and GISTIC copy number datasets of Breast cancer TCGA samples (1078 tumors, 17088 genes, 50 Hallmark genesets) or all TCGA samples (10459 tumors, 17446 genes, 50 Hallmark gene sets).
